# Rubisco adaptation is more limited by phylogenetic constraint than by catalytic trade-off

**DOI:** 10.1101/2020.09.15.298075

**Authors:** Jacques W. Bouvier, David M. Emms, Timothy Rhodes, Jai S. Bolton, Amelia Brasnett, Alice Eddershaw, Jochem R. Nielsen, Anastasia Unitt, Spencer M. Whitney, Steven Kelly

## Abstract

Rubisco assimilates CO_2_ to form the sugars that fuel life on earth. Correlations between rubisco kinetic traits across species have led to the proposition that rubisco adaptation is highly constrained by catalytic trade-offs. However, these analyses did not consider the phylogenetic context of the enzymes that were analysed. Thus, it is possible that the correlations observed were an artefact of the presence of phylogenetic signal in rubisco kinetics and the phylogenetic relationship between the species that were sampled. Here, we conducted a phylogenetically-resolved analysis of rubisco kinetics and show that there is a significant phylogenetic signal in rubisco kinetic traits. We re-evaluated the extent of catalytic trade-offs accounting for this phylogenetic signal and found that all were attenuated. Following phylogenetic correction, the largest catalytic trade-offs were observed between the Michaelis constant for CO_2_ and carboxylase turnover (∼21-37 %), and between the Michaelis constants for CO_2_ and O_2_ (∼9-19 %), respectively. All other catalytic trade-offs were substantially attenuated such that they were marginal (<9 %) or non-significant. This phylogenetically resolved analysis of rubisco kinetic evolution also identified kinetic changes that occur concomitant with the evolution of C_4_ photosynthesis. Finally, we show that phylogenetic constraints (most likely caused by a slow rate of molecular evolution) have played a larger role than catalytic trade-offs in limiting the evolution of rubisco kinetics. Thus, although there is strong evidence for some catalytic trade-offs, rubisco adaptation has been more limited by phylogenetic constraint than by the combined action of all such trade-offs.

## Main text

## Introduction

The vast majority of organic carbon on Earth entered the biosphere via the catalytic pocket of rubisco (ribulose-1,5-bisphosphate carboxylase/oxygenase) (Beer *et al*., 2010). Whilst there are several metabolic contexts in which this rubisco-mediated reaction can occur, the most important of these in terms of global net primary production is photosynthesis (Andersson and Backlund, 2008). Here, rubisco catalyses the initial step of the Calvin-Benson-Bassham reductive pentose phosphate pathway, catalysing the fixation of CO_2_ onto the acceptor molecule ribulose 1,5-bisphosphate (RuBP) to ultimately synthesise sugars. There is a diverse array of rubisco forms found across the tree of life. Plants and some bacteria contain Form I rubisco which is composed of both large (RbcL) and small (RbcS) subunits (Schneider, Lindqvist and Brändén, 1992; Tabita *et al*., 2008), while Form II, III and IV rubisco found in other lineages are comprised of just the large subunit (Tabita *et al*., 2008; Banda *et al*., 2020). Although Form I rubisco contains two subunits, only the large subunit is essential for catalysis (Andrews, 1988; Lee and Tabita, 1990; Whitney, Houtz and Alonso, 2011), while the small subunit has an indirect effect on catalytic properties and activity (Andrews, 1988; Lee and Tabita, 1990; Lee, Berka and Tabita, 1991; Read and Tabita, 1992b, 1992a; Spreitzer, Peddi and Satagopan, 2005; Ishikawa *et al*., 2011; Joshi *et al*., 2015; Fukayama *et al*., 2019; Martin-Avila *et al*., 2020).

Given that rubisco is the entry point for carbon into the global food chain it is perhaps unsurprising that it is the most abundant enzyme on Earth (Ellis, 1979) with a global mass of ∼0.7 gigatons (Bar-On and Milo, 2019). However, this abundance is in part due to the inefficiency of rubisco as a catalyst. Specifically, rubisco has a low rate of CO_2_ assimilation (Tcherkez, Farquhar and Andrews, 2006; Savir *et al*., 2010) and is poorly able to discriminate CO_2_ and O_2_ (Ogren and Bowes, 1971) causing it to catalyse both a carboxylation and an oxygenation reaction (Bowes, Ogren and Hageman, 1971; Chollet, 1977; Sharkey, 2020). Rubisco-mediated oxygenation of RuBP results in the production of 2-phosphoglycolate, which must then be metabolised to recover carbon and avoid depletion of metabolite pools (Eckardt, 2005; Sharwood, 2017). In plants this carbon scavenging process is known as photorespiration, and consumes ATP and reducing power and also liberates ammonia which must be re-assimilated (Peterhansel *et al*., 2010). Although the oxygenation reaction catalysed by rubisco is not thought to be deleterious in the anoxic environment prevalent when the enzyme first evolved (Nisbet *et al*., 2007; Erb and Zarzycki, 2018), under current atmospheric conditions it can comprise a quarter of all rubisco reactions in terrestrial plants (Ehleringer *et al*., 1991). Thus, despite oxygenation serving a number of beneficial functions (Busch, 2020), at its current rate it represents a substantial metabolic burden reducing the productivity of some plants by up to 50 % (Ogren, 1984; Bauwe *et al*., 2012).

Given the high energic cost incurred by the rubisco oxygenation reaction, a number of photoautotrophic organisms have evolved mechanisms to reduce the frequency of its occurrence (Flamholz and Shih, 2020). Collectively referred to as CO_2_-concentrating mechanisms, these function to increase the concentration of CO_2_ relative to O_2_ in the vicinity of rubisco and thus increase the relative frequency of carboxylation reactions (Meyer and Griffiths, 2013; Flamholz and Shih, 2020). The observation that evolution has resulted in an array of CO_2_-concentrating mechanisms, rather than improve the CO_2_ specificity of rubisco, has led many to question whether altering the kinetics of the enzyme is possible (Ogren, 1984; Parry *et al*., 2007, 2013; Whitney, Houtz and Alonso, 2011; Sharwood, Ghannoum and Whitney, 2016; Araújo *et al*., 2017; Sharwood, 2017; Sharkey, 2020). This proposition that rubisco specificity cannot be improved was supported by observations that the oxygenase and carboxylase activities of rubisco appear to be tightly linked (Badger and Lorimer, 1976; Chollet and Anderson, 1976). Subsequently, multiple studies have supported this suggestion by reporting strong antagonistic relationships between rubisco specificity (*S*_C/O_), carboxylase turnover (*k*_catC_) and the Michaelis constant (i.e., an inverse measure of substrate affinity for an enzyme) for CO_2_ (*K*_C_), as well as between *K*_C_ and the Michaelis constant for O_2_ (*K*_O_) (Tcherkez, Farquhar and Andrews, 2006; Savir *et al*., 2010). Collectively, these studies have led to the hypothesis that severe kinetic trait trade-offs hamstring the inherent efficiency by which the enzyme can catalyse CO_2_ fixation, and that contemporary rubisco are near perfectly adapted within this heavily constrained catalytic landscape (Tcherkez, Farquhar and Andrews, 2006; Savir *et al*., 2010). However, new evidence has begun to question this paradigm of rubisco evolution. First, recent analyses of the correlative nature of rubisco kinetics has demonstrated that associations between kinetic traits are weakened when a large number of species are considered (Flamholz *et al*., 2019; Iñiguez *et al*., 2020). Furthermore, engineering efforts to alter rubisco kinetics have produced enzyme variants that deviate from proposed catalytic trade-offs between *S*_C/O_, *k*_catC_ and *K*_C_ (Wilson *et al*., 2018; Zhou and Whitney, 2019). Finally, an updated examination of rubisco kinetics in the context of other enzymes has shown that it is not as inefficient a catalyst as often assumed (Bathellier *et al*., 2018). Thus, together these results indicate that rubisco kinetic traits are perhaps not as inextricably linked as originally thought, and suggest that there is scope for increasing the catalytic efficiency of the enzyme as has happened in nature for rubisco in some red algae (Andersson and Backlund, 2008; Gunn *et al*., 2020).

Although the kinetic traits of rubisco appear to be correlated, there are flaws to inferring causality from this correlation. This is because previous analyses that have inferred correlations have assumed that measurements of rubisco kinetic traits in different species are independent (Tcherkez, Farquhar and Andrews, 2006; Savir *et al*., 2010; Flamholz *et al*., 2019; Iñiguez *et al*., 2020). However, this assumption has never formally been tested and is unlikely to be true because rubisco in all species are related to each other by descent from a single ancestral gene. This means that the gene sequences encoding the enzyme in closely related species are more similar than the gene sequences from species that are more distantly related, a feature of rubisco which has long been exploited in systematics and evolutionary analyses to serve as an accurate proxy for the phylogenetic relationship between species (Gielly and Taberlet, 1994; APG, 1998, 2016). As sequence variation determines kinetic variation, closely related enzymes would be expected to also have similar kinetics, with the extent of this similarity being dependent on the underlying tree describing the relationship between species. This phenomenon, which is known as phylogenetic signal, can cause spurious correlations in measured trait values between species unless the structure of the phylogenetic tree is taken into consideration (Felsenstein, 1985; Grafen, 1989; Pagel and Harvey, 1989; Garland, 2001). Thus, as previous analyses of rubisco kinetics have not assessed whether a phylogenetic signal exists in rubisco kinetic traits, nor accounted for any phylogenetic signal which may exist, it is possible that the observed catalytic trade-offs inferred from the presence of correlations are, either wholly or in part, an artefact caused by this phylogenetic signal.

Here, we assess the presence of a phylogenetic signal in rubisco kinetic traits to evaluate the extent to which rubisco kinetic evolution is constrained by both phylogenetic effects and catalytic trade-offs. We demonstrate that there is a significant phylogenetic signal in all rubisco kinetic traits. This means that the similarity of kinetic measurements between species varies as a function of their evolutionary distance, and thus kinetic measurements in different species are non-independent. When this phylogenetic signal is correctly accounted for by using phylogenetic least squares regression, we reveal that inferred catalytic trade-offs are weakened and that rubisco kinetic traits have been evolving largely independently of each other. Moreover, we find that phylogenetic constraints, most likely resulting a slow rate of molecular evolution, have constrained rubisco kinetic evolution to a greater extent than catalytic trade-offs. This new insight offers encouragement to efforts which aim to increase yields in food, fibre and fuel crops by engineering rubisco variants with increased catalytic efficiency.

## Results

### Rubisco kinetic data

A dataset comprising kinetic measurements for rubisco isolated from different photoautotrophs was obtained from (Flamholz *et al*., 2019). Measurements of specificity (*S*_C/O_) for CO_2_ relative to O_2_ (i.e., the overall carboxylation/oxygenation ratio of rubisco under defined concentrations of CO_2_ and O_2_ gases) in this dataset were normalised in order to overcome discrepancies between values determined using an oxygen electrode assay (Parry, Keys and Gutteridge, 1989) and high precision gas-phase-controlled 3H-RuBP-fixation assays (Kane et al., 1994) (see methods). To begin, the interrogation of this data was focussed on the angiosperms because this was the group with the largest and most complete set of kinetic measurements, and to minimize any impact of long-branch effects (Su and Townsend, 2015). It was also restricted to those species with measurements of *S*_C/O_, maximum carboxylase turnover rate per active site (*k*_catC_), and the Michaelis constant (i.e., the substrate concentration at half-saturated catalysed rate) for both CO_2_ (*K*_C_) and O_2_ (*K*_O_). The Michaelis constant for CO_2_ in 20.95 % O_2_ air (*K*_C_^air^) was also inferred as a function of both *K*_C_ and *K*_O_ (see methods). Of the 137 angiosperms that satisfied these filtration criteria, 19 also had measurements of the Michaelis constant for RuBP (*K*_RuBP_). From here on, these constants and rates are collectively termed kinetic traits, where *S*_C/O_, *k*_catC_, *K*_C_ and *K*_C_^air^ are referred to as carboxylase-related kinetic traits, and *K*_O_ as the oxygenase-related kinetic trait.

### A significant phylogenetic signal exists in rubisco carboxylase-related kinetic traits in angiosperms

Consistent with previous analyses (Flamholz *et al*., 2019), all kinetic traits were log transformed to ensure they conformed to the distribution assumptions of the statistical analyses herein. To assess whether rubisco in different angiosperms display similar kinetics as a consequence of their phylogenetic relationship, the kinetic traits were analyzed in the context of the phylogenetic tree by which the species are related (Figure 1). Here, all kinetic traits were subject to interrogation for a phylogenetic signal (Table 1) except for *K*_RuBP_, which was omitted owing to the limited number of measurements available for this trait. For these analyses, several statistical tools varying in their approach to phylogenetic signal detection were implemented and the presence or absence of a phylogenetic signal in each trait was judged by the majority result (i.e., the result of ≥ 3 out of 5 methods tested). Out of the methods utilized, Pagel’s lambda (Pagel, 1999) and Blomberg’s K and K* (Blomberg, Garland and Ives, 2003) analyse the distribution of trait values in extant species using an explicit Brownian motion model of trait evolution in which the traits evolve stochastically on the underlying phylogenetic tree at a uniform rate and independently among branches. In contrast, Moran’s I (Gittleman and Kot, 1990) and Abouheif’s Cmean (Abouheif, 1999) do not invoke any specific aspect of evolutionary theory, but instead test for a phylogenetic signal by assessing the correlation of trait values across evolutionary distance on the species tree using the concept of autocorrelation adopted from the field of spatial statistics (Cheverud, Dow and Leutenegger, 1985, 1986). For further discussion of the differences between these phylogenetic signal detection methods see (Münkemüller *et al*., 2012).

**Figure 1.**
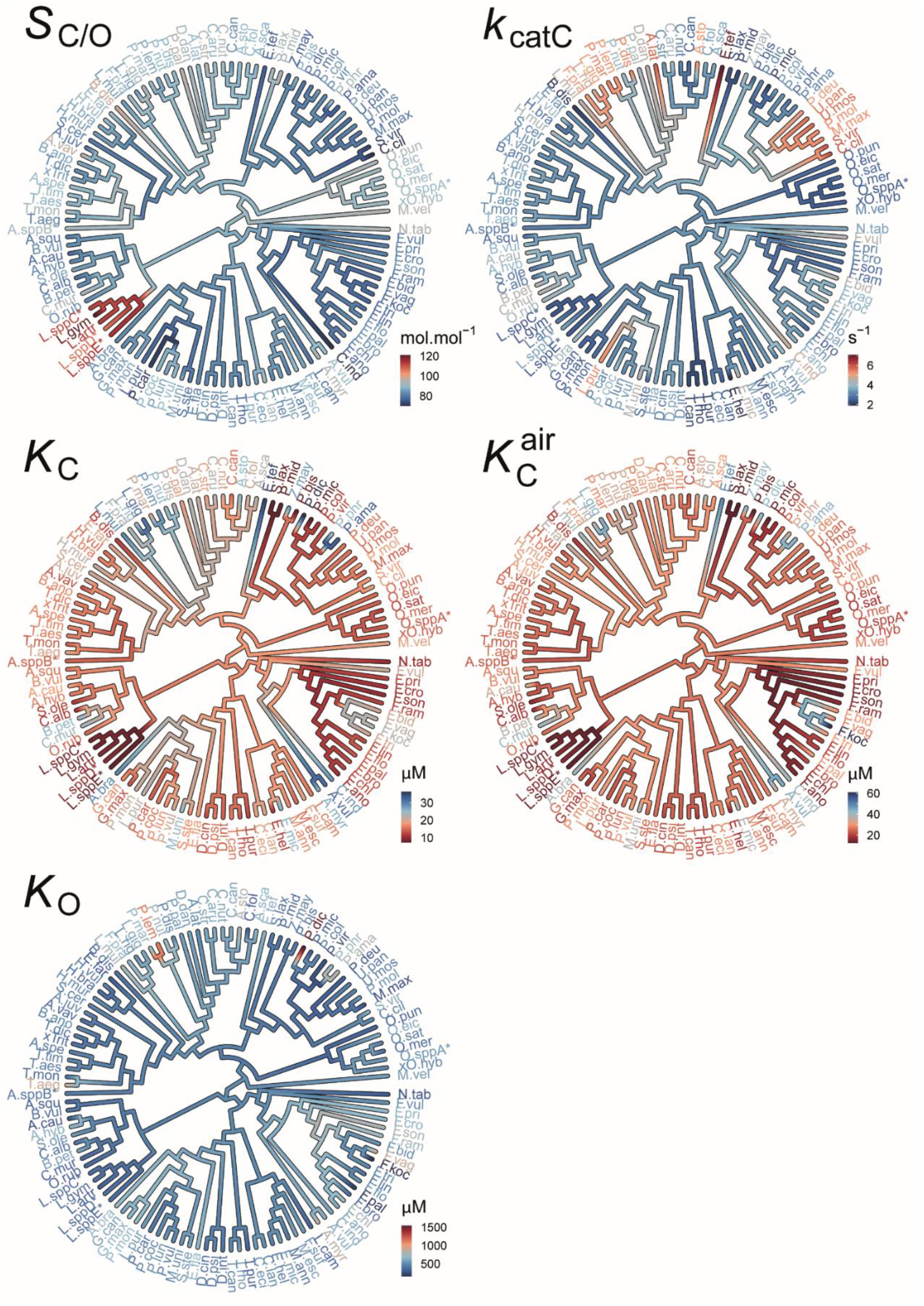
The evolution of rubisco kinetic traits in angiosperms. Phylogenetic tree of angiosperm species depicting the kinetic trait values in of the species used in this dataset and the maximum likelihood inferred ancestral kinetic traits for internal branches on the tree. Scale bars for colour schemes are presented next to each tree. Species names have been abbreviated for legibility and are provided in full in Supplementary File 1, Table S4. *S*_C/O_: specificity. *k*_catC_: carboxylase turnover per site. *K*_C_: the Michaelis constant for the CO_2_. *K*_C_^air^ the inferred Michaelis constant for CO_2_ in 20.95 % O_2_ air. *K*_O_: the Michaelis constant for O_2_.

**Table 1.** The phylogenetic signal strength and associated significance level in rubisco kinetic traits in angiosperms using five different signal detection methods. Statistics are rounded to 3 decimal places and significance values are represented as α levels, where; α = 0.001 if *p* < 0.001, α = 0.01 if 0.001 < *p* < 0.01, α = 0.05 if 0.01 < *p* < 0.05, and α = ns if *p* > 0.05.

| Kinetic Trait | C mean |  | I |  | K |  | K * |  | Lambda |  |
| --- | --- | --- | --- | --- | --- | --- | --- | --- | --- | --- |
| | Stat | $\alpha$ | Stat | $\alpha$ | Stat | $\alpha$ | Stat | $\alpha$ | Stat | $\alpha$ |
| **S<sub>C/O</sub>** | 0.516 | 0.001 | 0.425 | 0.001 | 0.001 | ns | 0.001 | ns | 0.879 | 0.001 |
| **k<sub>catC</sub>** | 0.350 | 0.001 | 0.224 | 0.001 | 0.003 | 0.001 | 0.003 | 0.001 | 0.968 | 0.001 |
| **K<sub>C</sub>** | 0.282 | 0.001 | 0.248 | 0.001 | 0.001 | 0.01 | 0.002 | 0.01 | 0.902 | 0.001 |
| **K<sub>C</sub><sup>air</sup>** | 0.234 | 0.001 | 0.242 | 0.001 | 0.001 | 0.05 | 0.001 | 0.05 | 0.392 | ns |
| **K<sub>O</sub>** | 0.032 | ns | -0.070 | ns | 0 | ns | 0 | ns | 0 | ns |

Irrespective of the methodological approach used for inference, a significant phylogenetic signal was observed in all carboxylase-related kinetic traits of *S*_C/O,_ *k*_catC,_ *K*_C_, and *K*_C_^air^ in angiosperms (Table 1; Figure 1). However, the strength of this signal varied across the different methods (Table 1). In contrast, a phylogenetic signal was not detected in angiosperms for the oxygenase-related kinetic trait *K*_O_ (Table 1; Figure 1). These measurements of phylogenetic signal were demonstrated to not suffer from overfitting due to the use of the *rbcL* gene to infer the phylogenetic tree (Supplementary File 1, Figure S1; Figure S2; and Table S1). Overall, this means that the similarity in carboxylase-related (but not oxygenase-related) kinetic traits in different angiosperms is dependent on their phylogenetic relationship. Therefore, conventional approaches to measure correlations that assume independence between observations of carboxylase-related kinetic trait values are incorrect, and correlation coefficients computed using such approaches have likely been over-estimated (Felsenstein, 1985; Grafen, 1989; Pagel and Harvey, 1989; Gittleman and Kot, 1990; Abouheif, 1999; Pagel, 1999; Garland, 2001; Blomberg, Garland and Ives, 2003).

### Significant changes in rubisco kinetics occur during the evolution of C_4_ photosynthesis

Inspection of the data identified several dependencies in rubisco kinetic traits between C_3_ and C_4_ plants (Figure 2). Specifically, the mean of the distribution of rubisco *S*_C/O_ values in C_4_ species (mean *S*_C/O_ = 78.7 mol.mol^-1^) was lower than that observed for rubisco in C_3_ species (mean *S*_C/O_ = 89.9 mol.mol^-1^) (Figure 2; p < 0.001, t-test). Conversely, the mean of the distribution of rubisco *k*_catC_ values was higher in C_4_ species (mean *k*_catC_ = 4.2 s^-1^) than in C_3_ species (mean *k*_catC_ = 3.2 s^-1^) (Figure 2; p < 0.001, t-test). The means of the distributions of both *K*_C_ and *K*_C_^air^ were also found to be higher in C_4_ species (mean *K*_C_ = 19.0 µM, mean *K*_C_^air^ = 29.9 µM) than in C_3_ species (mean *K*_C_ = 15.4 µM, mean *K*_C_^air^ = 23.6 µM) (Figure 2; p < 0.05 and p < 0.05, t-test, respectively). In contrast, no significant difference was observed in *K*_O_ between C_3_ species (mean *K*_O_ = 481.0 µM) and C_4_ species (mean *K*_O_ = 466.7 µM) (Figure 2; p > 0.05, t-test). However, variation in *K*_O_ was found to be considerably greater in C_4_ species (95 % CI = [379.1, 574.6]) than in C_3_ species (95 % CI = [457.1, 506.0]) (p < 0.01; Levene test). Further, although the restricted number of *K*_RuBP_ measurements did not allow statistical differences to be assessed between photosynthetic groups, the distribution of this trait appeared to show higher variability in C_4_ species, similar to that observed for *K*_O_. Owing to limited kinetic measurements for rubisco in C_3_-C_4_ intermediate and C_4_-like species which respectively represent early and late transition states along the evolutionary continuum from C_3_ to C_4_ photosynthesis, it was not possible to assess changes in rubisco kinetics in these plants relative to the ancestral C_3_ and derived C_4_ photosynthetic types. Nevertheless, trait values of rubisco *S*_C/O_ in both evolutionary intermediate C_3_-C_4_ and C_4_-like states appear to closely resemble the distribution observed in C_4_ species (Figure 2), thus indicating adaptation of this trait potentially occurred early during the evolution of C_4_ photosynthesis. All of the significant differences between C_3_ and C_4_ plants reported above were robust to correction for the measured phylogenetic signal (Supplementary File 1, Table S2). Collectively, these data demonstrate that there are adaptations in rubisco kinetics that are associated with the evolution of C_4_ photosynthesis, such that the emergence of the C_4_ carbon concentrating mechanism is accompanied by a decreased specificity and CO_2_ affinity, and an increased carboxylase turnover.

**Figure 2.**
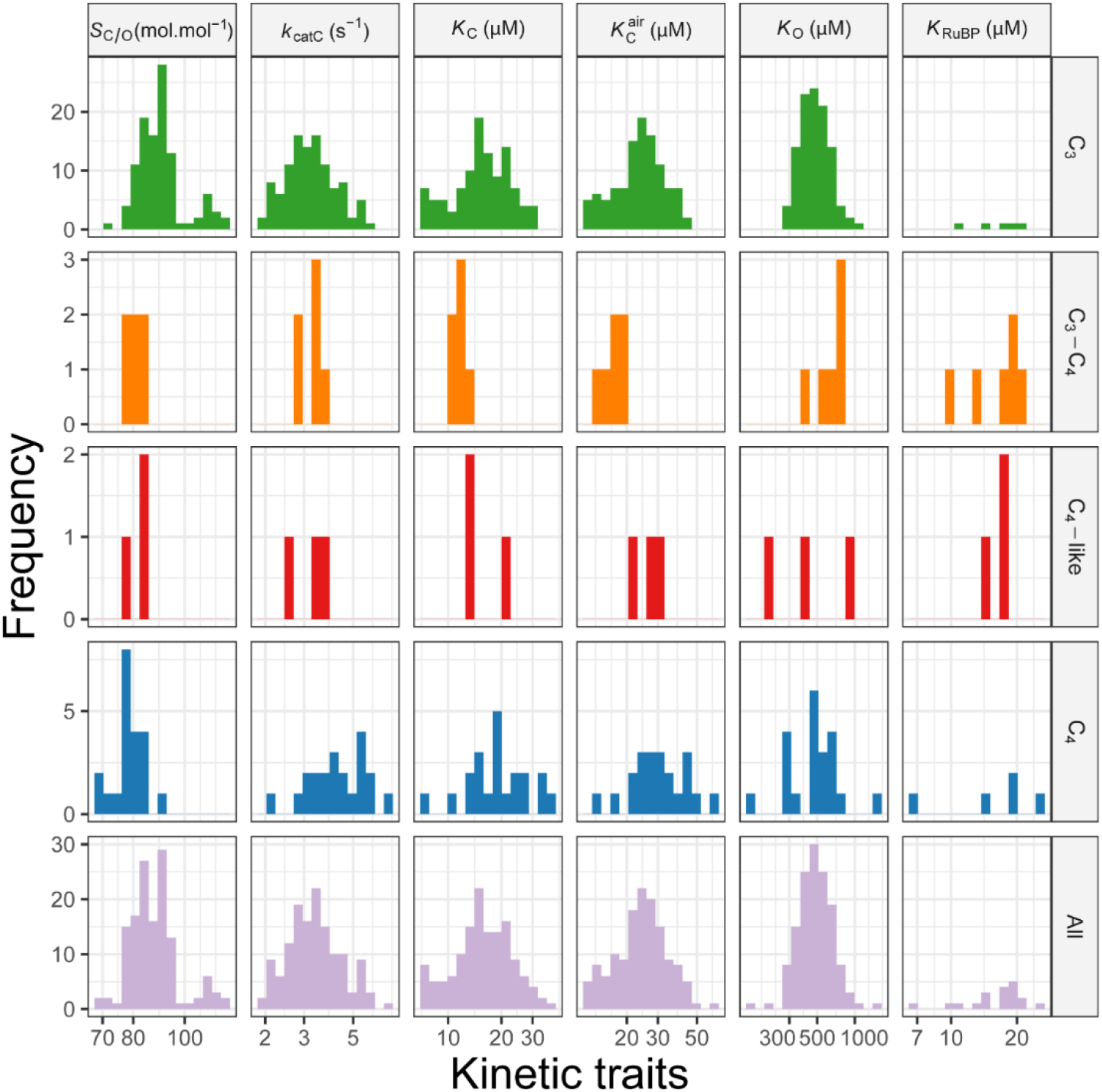
The distributions of values for rubisco kinetic traits in angiosperms. Species are grouped by their photosynthetic types (rows). *S*_C/O_: specificity. *k*_catC_: carboxylase turnover per site. *K*_C_: the Michaelis constant for the CO_2_. *K*_C_^air^ the inferred Michaelis constant for CO_2_ in 20.95 % O_2_ air. *K*_O_: the Michaelis constant for O_2_. *K*_RuBP_: the Michaelis constant for ribulose 1,5-bisphosphate. Plants have been classified as those which perform C_3_ photosynthesis (C_3_; *n* = 107), C_4_ photosynthesis (C_4_; *n* = 6), C_3_-C_4_ intermediate (C_3_-C_4_; *n* = 3), C_4_-like (C_4_-like; *n* = 21). The X axis for all plots is on a log scale, where respective units are shown in column labels. The raw dataset used can be found in Supplemental File 2.

### A significant phylogenetic signal exists in rubisco K_O_ in C_3_ plants

Based on the positions of C_3_-C_4_ intermediate, C_4_-like, and C_4_ species in the phylogenetic tree (Supplementary File 1, Figure S1), multiple independent evolutions toward C_4_ photosynthesis are present in the dataset. Furthermore, given that transition to C_4_ photosynthesis is found above to be associated with adaptive changes in rubisco kinetic traits including a reduction in *S*_C/O_, an increase in *k*_catC_, *K*_C_ and *K*_C_^air^, as well as increased variability in *K*_O_ (Figure 1; Supplementary File 1, Table S2), it was hypothesised that a failure to account for kinetic differences associated with photosynthetic type may have confounded estimations of the phylogenetic signal. For example, kinetic modifications associated with the evolution of C_4_ photosynthesis may cause larger differences in rubisco kinetics among closely related C_3_ and C_4_ species than expected based on evolutionary distance alone. Similarly, the independent evolution of C_4_ photosynthesis in distantly related plant lineages could also cause evolutionarily distant species to have similar kinetic trait values by convergence. To evaluate the extent by which these respective issues may have affected quantification of the phylogenetic signal, the above analyses were repeated using only the C_3_ angiosperm species present (i.e., with C_3_-C_4_ intermediate, C_4_-like, and C_4_ species removed). In general, estimates of the phylogenetic signal in the carboxylase-related kinetic traits in C_3_ species (Table 2) agreed with those observed when all angiosperms were considered on the phylogenetic tree (Table 1). Specifically, a phylogenetic signal of similar strength and significance was observed in *S*_C/O_, *k*_catC_ and *K*_C_ for each of the detection methods across both sets of analyses (Table 1 and 2). In addition, the discrepancies in signal strength between the statistical methods previously observed (Table 1) were recapitulated in the analysis using only C_3_ species (Table 2), thus indicating that these differences are not caused by a failure to control for photosynthetic type, but instead more likely represent distinctions in the assumptions and aspects of the phylogenetic signal measured by each test (Hardy and Pavoine, 2012; Münkemüller *et al*., 2012). In summary, there is a statistically significant phylogenetic signal in rubisco specificity, carboxylase turnover and the Michaelis constant for CO_2_ in angiosperms that is independent of photosynthetic type.

**Table 2.** The phylogenetic signal strength and associated significance level in rubisco kinetic traits in C_3_ species using five different signal detection methods. Statistics are rounded to 3 decimal places, and significance values are represented as α levels, where; α = 0.001 if *p* < 0.001, α = 0.01 if 0.001 < *p* < 0.01, α = 0.05 if 0.01 < *p* < 0.05, and α = ns if *p* > 0.05.

| Kinetic Trait | C mean |  | I |  | K |  | K * |  | Lambda |  |
| --- | --- | --- | --- | --- | --- | --- | --- | --- | --- | --- |
| | Stat | $\alpha$ | Stat | $\alpha$ | Stat | $\alpha$ | Stat | $\alpha$ | Stat | $\alpha$ |
| **S<sub>C/O</sub>** | 0.533 | 0.001 | 0.453 | 0.001 | 0 | ns | 0.001 | ns | 0.814 | 0.001 |
| **k<sub>catC</sub>** | 0.387 | 0.001 | 0.234 | 0.001 | 0.002 | 0.01 | 0.002 | 0.01 | 0.913 | 0.001 |
| **K<sub>C</sub>** | 0.449 | 0.001 | 0.341 | 0.001 | 0.001 | 0.05 | 0.001 | 0.05 | 0.948 | 0.001 |
| **K<sub>C</sub><sup>air</sup>** | 0.398 | 0.001 | 0.317 | 0.001 | 0.001 | 0.05 | 0.002 | 0.01 | 0.947 | 0.001 |
| **K<sub>O</sub>** | 0.279 | 0.001 | 0.167 | 0.01 | 0 | ns | 0 | ns | 0.743 | 0.001 |

In contrast to the analysis of all angiosperms (Table 1), a significant signal was observed in *K*_O_ when only C_3_ angiosperms were considered (Table 2). Thus, both the oxygenase-related and carboxylase-related traits of rubisco have evolved in a tree-like manner in C_3_ angiosperms. Furthermore, unlike the other carboxylase-related kinetic traits, the phylogenetic signal in *K*_C_^air^ was found to increase in strength when the analysis is restricted to C_3_ angiosperms. This result is a corollary of the fact that *K*_C_^air^ is computed here from both *K*_C_ and *K*_O_. Thus, all kinetic traits of rubisco have a significant phylogenetic signal in C_3_ angiosperms.

### Correlations between kinetic traits are weak in angiosperms and further relaxed after correcting for a phylogenetic signal

Given the finding that rubisco kinetic traits exhibit a significant phylogenetic signal (Table 1; Figure 2), it is possible that previously reported correlations between rubisco kinetic traits (Tcherkez, Farquhar and Andrews, 2006; Savir *et al*., 2010; Flamholz *et al*., 2019; Iñiguez *et al*., 2020) are an artefact of this signal. This is because prior analyses which did not account for the inherent phylogenetic structure (and non-independence of) this data (Figure 3A) may have suffered from over-estimated correlation coefficients due to this underlying structure. In order to evaluate the severity by which this signal may have influenced previous results (Tcherkez, Farquhar and Andrews, 2006; Savir *et al*., 2010; Flamholz *et al*., 2019; Iñiguez *et al*., 2020), the correlations observed in the kinetic trait data using both phylogenetic and non-phylogenetic regression methods were compared (Figure 3B and 3C).

**Figure 3.**
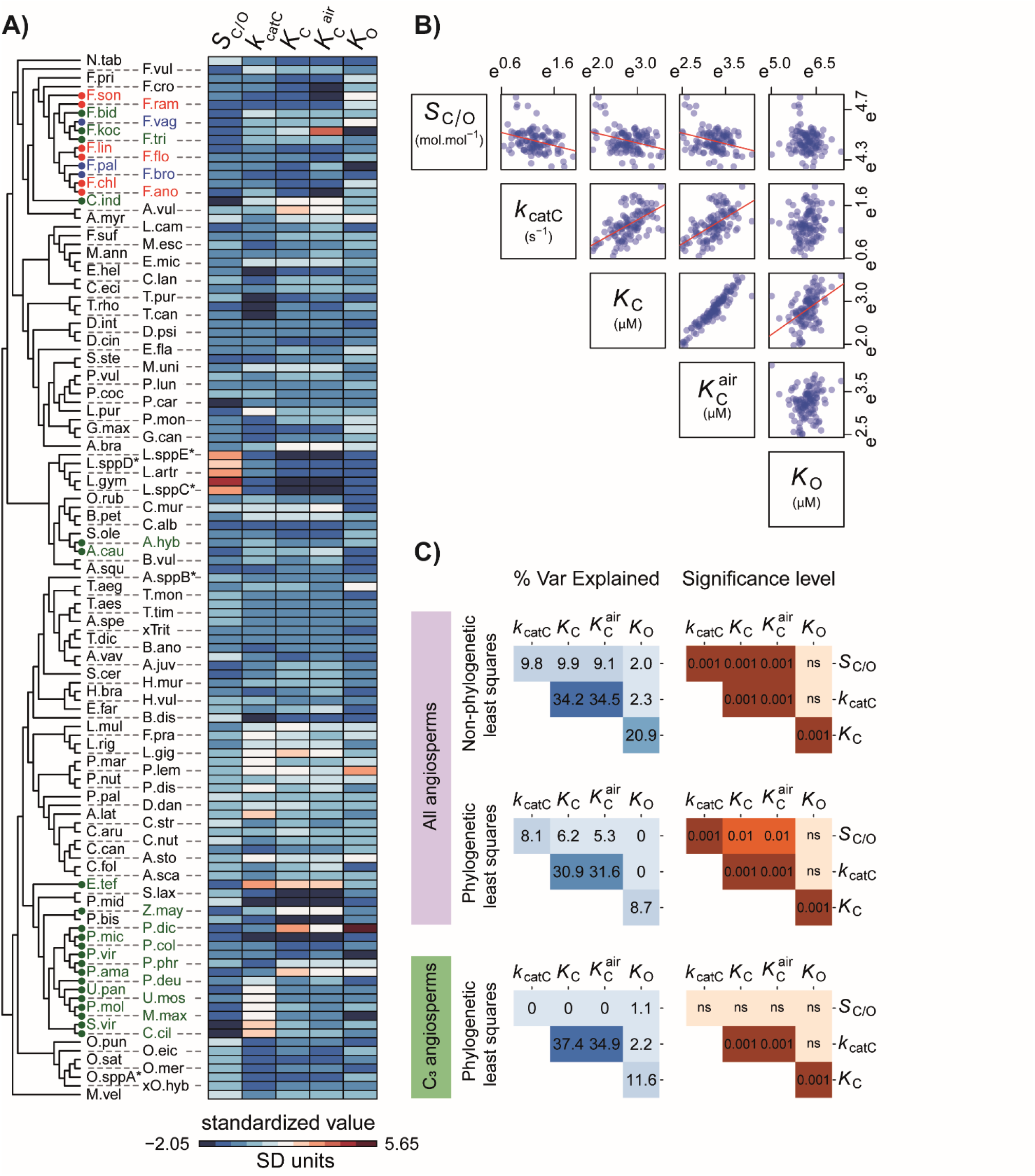
The correlations between rubisco kinetic traits in angiosperms. **A**) Heatmap depicting the normalised variation in kinetic traits across the species set used in this study (± S.D. away from each respective kinetic trait mean). Species labels on the tree are colour coded by photosynthetic type (C_3_: black, C_3_-C_4_ intermediates: red, C_4_-like: blue, and C_4_: green), and have been abbreviated for legibility (refer to Supplemental File 1, Table S4). **B)** Trends in relationships between all pairwise combinations of log transformed rubisco kinetic traits shown in A. **C**) Pairwise correlation coefficients (percent variance explained) and associated *p*-values between different rubisco kinetic traits assessed using non-phylogenetic least squares regression models or phylogenetic least squares regression models. Phylogenetic least squares regression was fit to both the complete set of angiosperms in the dataset and the subset which perform C_3_ photosynthesis. Significance values are represented as α levels, where; α = 0.001 if *p* < 0.001, α = 0.01 if 0.001 < *p* < 0.01, α = 0.05 if 0.01 < *p* < 0.05, and α = ns if *p* > 0.05.

Using a standard non-phylogenetic approach, the relationships between kinetic traits of rubisco were consistent in both linear and least squares regression models (Supplementary File 1, Figure S3A and S3B). The direction of power-law relationships observed (Figure 3B) match those previously reported (Flamholz *et al*., 2019). Specifically, significant positive correlations were found between *k*_catC_ and both *K*_C_ and *K* ^air^ (Figure 3B and 3C). A significant positive correlation was also observed between the respective Michaelis constants for both CO_2_ and O_2_ substrates, *K*_C_ and *K*_O_ (Figure 3B and 3C). In addition, significant inverse power-law correlations were observed between *S*_C/O_ and all other carboxylase-related kinetic traits investigated, including *k*_catC_, *K*_C_ and *K*_C_^air^ (Figure 3B and 3C). In contrast, *K*_O_ did not co-vary with either *S*_C/O_ or *k*_catC_ (Figure 3B and 3C), whilst *K*_RuBP_ did not appear to be tightly linked to any kinetic trait from the limited number of observations that are available (Supplementary File 1, Figure S3A). Thus, all pairwise relationships between the carboxylase-related kinetic traits *S*_C/O_, *k*_catC_ and either *K*_C_ or *K*_C_^air^ were significant, whilst the oxygenase-related trait *K*_O_ was only correlated with *K*_C_.

Although kinetic trade-offs inferred using non-phylogenetic methods were concordant in direction with those previously described (Flamholz *et al*., 2019), they were substantially reduced in magnitude when the analysis was focussed solely on the angiosperms. This reduction in magnitude of correlation when taxonomic groups are removed is strongly indicative of phylogenetic signal in the wider dataset and is analysed further detail in a subsequent results section. Within angiosperms, the strength of the correlation between *S*_C/O_ and *K*_C_ (9.9 % variance explained; Figure 3C) is attenuated by 77 % when compared to that previously reported using a larger range of rubisco variants (43.6 % variance explained (Flamholz *et al*., 2019)). Moreover, a 69 % reduction was found in the dependency between *S*_C/O_ and *k*_catC_ in angiosperms (9.8 % variance explained; Figure 3C) in comparison to that reported based on the larger range of species (31.4 % variance explained (Flamholz *et al*., 2019)), with the antagonistic correlation observed between *K*_C_ and *K*_O_ (20.9 % variance explained; Figure 3C) also weakened by 33 % relative to previous reports (31.4 % variance explained (Flamholz *et al*., 2019)). In contrast, the dependency between *K*_C_ and *k*_catC_ was 49 % stronger when only angiosperms are assessed, increasing from 23.0 % (Flamholz *et al*., 2019) to 34.2 % in this study (Figure 3C). Therefore overall, even in the absence of correctly accounting for the phylogenetic relationship between rubisco, the apparent catalytic trade-offs observed in angiosperms are weaker than previously thought (Flamholz *et al*., 2019; Iñiguez *et al*., 2020).

Given that a significant phylogenetic signal is present in rubisco kinetic traits in angiosperms (Table 1 and 2), a phylogenetic generalized least squares regression analysis (Felsenstein, 1985) was conducted to estimate the magnitude of the catalytic trade-offs accounting for the inherent structure of the data. In comparison to phylogeny-unaware correlations, the phylogenetic regression resulted in a reduction in the majority of kinetic trade-offs (Figure 3C). The largest reduction observed was for the correlation between the Michaelis constants *K*_C_ and *K*_O._ Here, the correlation was reduced by 58 % (variance explained = 8.7 %) relative to methods which do not correctly account for the non-independence of these measurements (variance explained = 20.9 %; Figure 3C). An analogous weak correlation was also observed when the phylogenetic analyses were limited to C_3_ species (variance explained = 11.6 %; Figure 3C). Thus, changes in *K*_C_ have occurred independently of any change on *K*_O_ in rubisco since the diversification of the angiosperms.

Phylogenetic correction also resulted in less substantial reductions in the correlation between *S*_C/O_ and each of the other carboxylase-related traits (Figure 3C). Here the dependency between *S*_C/O_ and *k*_catC_ was reduced by 18 % from 9.8 to 8.1 %, whilst the dependency between *S*_C/O_ and both *K*_C_ and *K*_C_^air^ were reduced by 37 % and 42 % from 9.9 to 6.2 %, and from 9.1 to 5.3 %, respectively (Figure 3C). Furthermore, these correlations were not significant when considering only C_3_ species (Figure 3C). Thus, during the evolution of rubisco in angiosperms, changes to specificity have occurred with little or no effect on other carboxylase-related kinetic traits, and *vice versa*.

In contrast, the strength of the correlation between *k*_catC_ and either *K*_C_ or *K*_C_^air^ was robust to phylogenetic correction. Specifically, the dependency between *k*_catC_ and *K*_C_ only decreased by 10 % from 34.2 to 30.9 %, and the dependency between *k*_catC_ and *K*_C_^air^ decreased by only 8 % from 34.5 to 31.6 %; Figure 3C). Further, the phylogenetically corrected correlations between these kinetic traits were of a similar magnitude when only C_3_ species were considered (37.4 and 34.9 % respectively; Figure 3C). Thus, as rubisco kinetics have evolved in angiosperms, there has been a trade-off between CO_2_ affinity and carboxylase turnover such that any change in one kinetic trait caused a partial change in the other, though with little impact on any further rubisco kinetic traits.

### The evolution of rubisco kinetics is more limited by phylogenetic constraints than by catalytic trade-offs in angiosperms

As rubisco kinetic traits contain a phylogenetic signal in angiosperms (Table 1 and 2), we sought to determine the extent to which the phylogenetic signal was caused by phylogenetic constraint. Here, phylogenetic constraint is considered to comprise all constraints which are embedded within the structure of the phylogenetic tree, that are independent of the kinetic constraints, and collectively act to impede the adaptive evolution of rubisco kinetics. For example, such phylogenetic constraints include processes which lead to neutral evolution (Felsenstein, 1985) or evolutionary stasis (Prinzing *et al*., 2001; Ackerly, 2004; Cavender-Bares *et al*., 2004; Moles *et al*., 2005; Swenson and Enquist, 2007) of the trait in question (Ives, 2019; Nevo *et al*., 2020). In order to assess the relative strength of such phylogenetic constraints on rubisco kinetics, the variance in kinetic traits partitioned between phylogenetic effects (i.e., the explanatory power of the phylogenetic tree in the goodness-of-fit model and a measure of phylogenetic constraint) and non-phylogenetic effects (i.e., the remaining explanatory power of the regression model, accounted for by the sum of all other constraints such as random effects and all kinetic trait trade-offs) was quantified. This analysis revealed that phylogenetic constraints explained a significant proportion of variation in carboxylase-related kinetic traits across all angiosperms (Figure 4A), and all kinetic traits in C_3_ angiosperms (Figure 4B). With one exception (i.e. the phylogenetic constraint on *K*_*O*_ in the larger species dataset) the magnitude of variation explained by phylogenetic constraints was similar or larger to the variation explained by the strongest trade-off observed between *k*_catC_ and *K*_C_ (Figure 4C and 4D). Consequently, across angiosperms the cumulative variance explained by phylogenetic constraints across all rubisco kinetic traits (29.5 %) was larger than the cumulative variance for all catalytic trade-offs combined (9.0 %). This effect was more pronounced for C_3_ angiosperms (cumulative variance for phylogenetic constraints = 43.4 %, cumulative variance for catalytic trade-offs = 8.2 %). Thus, during the radiation of angiosperms phylogenetic constraints have restricted the evolution of rubisco kinetics to a greater extent than all catalytic trade-offs combined.

**Figure 4.**
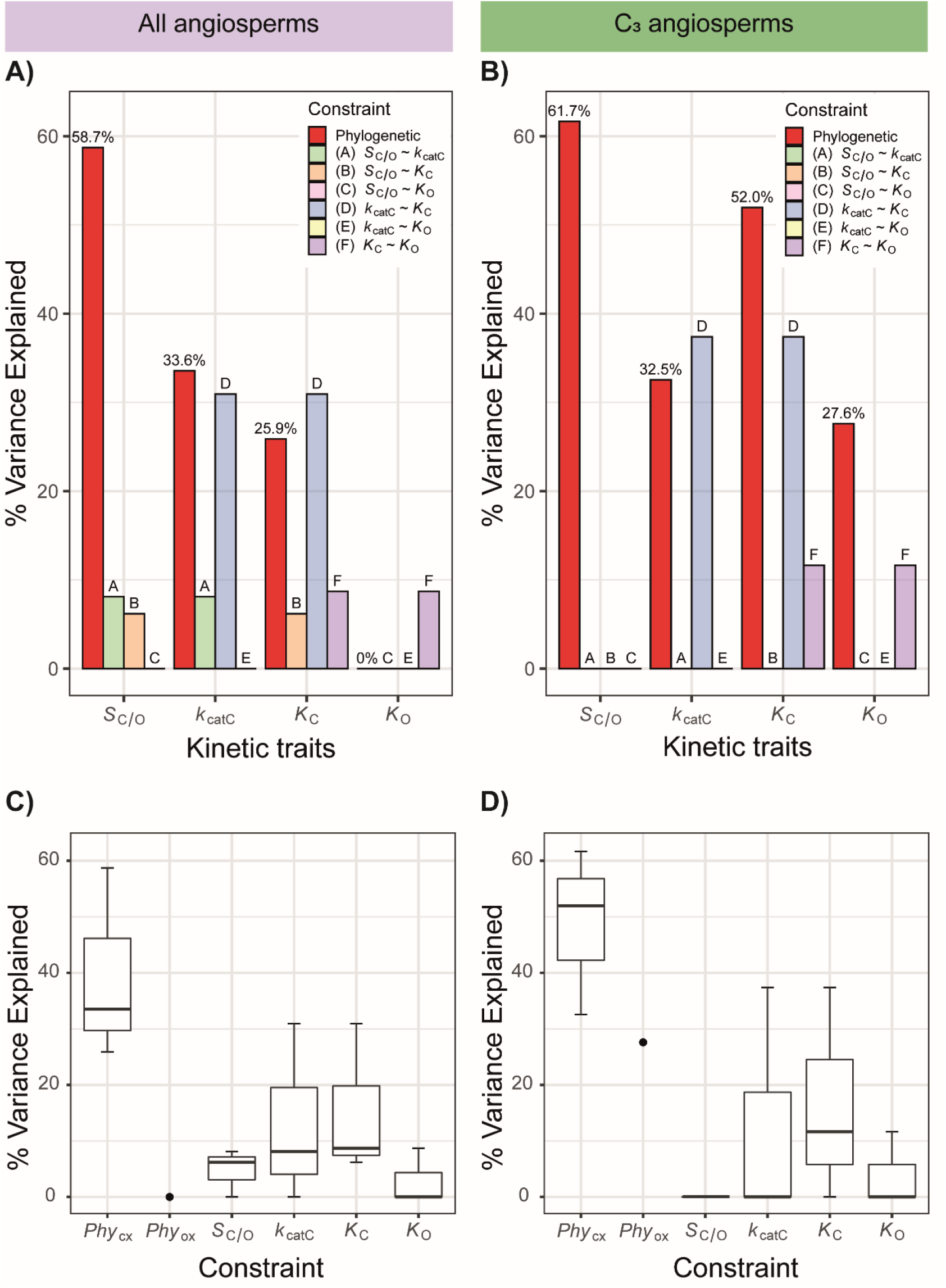
The constraints on rubisco kinetic adaptation in angiosperms. **A)** The variation (%) in rubisco kinetic traits across angiosperms that can be explained by phylogenetic constraint and each catalytic trade-off. **B)** As in A but for C_3_ angiosperms only. **C)** Boxplot of all variation explained in each kinetic trait by kinetic trait correlations in comparison to variation explained by phylogeny in angiosperms. The phylogenetic constraints on the carboxylase-related traits *Phy*_CX_ (includes *Phy*_*Sc/o*_, *Phy*_*Kcatc*_, and *Phy*_*Kc*_) and phylogenetic constraints on the oxygenase-related trait *Phy*_ox_ (includes *Phy*_*Ko*_ only) are presented separately. **D)** As in C but for C_3_ angiosperms only.

### Phylogenetic signal, weak kinetic trait correlations and strong phylogenetic constraint are features of rubisco in all photosynthetic organisms

Given the presence of phylogenetic signal and the impact of phylogenetic constraint on the evolution of rubisco kinetics in angiosperms, we sought to determine whether these findings were a unique feature of angiosperm rubisco or whether they were a more general phenomenon across the tree of life. To achieve this, the dataset was expanded to include all species for which both kinetic measurements and an *rbcL* gene sequence were available. Analogous to the analysis of angiosperms, a strong and statistically significant phylogenetic signal was observed in *S*_C/O_, *k*_catC_, *K*_C_ and *K*_C_^air^, but not in *K*_O_ across all photosynthetic organisms (Table 3). Similarly, a significant phylogenetic signal was also observed for *K*_O_ when C_3_-C_4_, C_4_-like and C_4_ angiosperms were omitted to control for the dependency in kinetic trait measurements on the tree associated with the convergent transition to C_4_ photosynthesis in angiosperms (Supplemental File 1, Table S3). Thus, there is a significant phylogenetic signal in rubisco kinetic traits in all photosynthetic organisms.

**Table 3.** The phylogenetic signal strength and associated significance level in rubisco kinetic traits across all studied photosynthetic organisms using five different signal detection methods. Statistics are rounded to 3 decimal places and significance values are represented as α levels, where; α = 0.001 if *p* < 0.001, α = 0.01 if 0.001 < *p* < 0.01, α = 0.05 if 0.01 < *p* < 0.05, and α = ns if *p* > 0.05.

| Kinetic<br>Trait | C mean |  | I |  | K |  | K * |  | Lambda |  |
| --- | --- | --- | --- | --- | --- | --- | --- | --- | --- | --- |
| | Stat | $\alpha$ | Stat | $\alpha$ | Stat | $\alpha$ | Stat | $\alpha$ | Stat | $\alpha$ |
| **S<sub>C/o</sub>** | 0.712 | 0.001 | 0.229 | 0.001 | 0.005 | 0.001 | 0.002 | 0.001 | 1.005 | 0.001 |
| **k<sub>catC</sub>** | 0.538 | 0.001 | 0.29 | 0.001 | 0.002 | 0.001 | 0.001 | 0.001 | 0.975 | 0.001 |
| **K<sub>C</sub>** | 0.705 | 0.001 | 0.277 | 0.001 | 0.002 | 0.001 | 0.001 | 0.001 | 0.948 | 0.001 |
| **K<sub>C</sub><sup>air</sup>** | 0.609 | 0.001 | 0.299 | 0.001 | 0.001 | 0.001 | 0.001 | 0.01 | 0.924 | 0.001 |
| **K<sub>O</sub>** | 0.170 | 0.001 | 0.004 | ns | 0 | ns | 0 | ns | 0.603 | 0.001 |

Analogous to above analyses, accounting for the phylogenetic tree caused a substantial attenuation in the kinetic trait correlations in all species (Figure 5A; Supplementary File 1, Figure S3C). Specifically, when correcting for the measured phylogenetic signal in kinetic traits, a partial correlation between *k*_catC_ and both *K*_C_ and *K*_C_^air^ was observed (variance explained = 21.3 % and 23.3 %, respectively; Figure 5A). Furthermore, a partial correlation was also measured between *K*_C_ and *K*_O_ (variance explained = 18.6 %; Figure 5A). However, correlations between all other pairwise combinations of kinetic traits were found to be either marginal, or not significant (Figure 5A). In addition, the dependency between *K*_C_ and *K*_O_ was attenuated to 13.4 % when the C_3_-C_4_, C_4_-like and C_4_ angiosperms were excluded from the analysis (Supplemental Supplementary File 1, Figure S5A). Evaluation of the phylogenetic constraints revealed that they explained a significant proportion of variation in the evolution in all rubisco kinetic traits (Figure 5B). Moreover, the phylogenetic constraints explained a larger proportion of kinetic trait variation than catalytic trade-offs (Figure 5C), such that the cumulate variation explained by phylogenetic constraints (56.1 %) was larger than the combined effect of all catalytic trade-offs (8.0 %). Analogous results were recovered when C_3_-C_4_, C_4_-like and C_4_ species were removed from the analysis (cumulative variance for phylogenetic constraints = 61.4 %, cumulative variance for catalytic trade-offs = 6.2 %; Supplemental Supplementary File 1, Figure S5B and S5C). Thus, phylogenetic constraints have been a critical limitation on rubisco adaptation in a diverse range of photoautotrophs and have presented a greater barrier to kinetic evolution compared to that imposed by all catalytic trade-offs combined.

**Figure 5.**
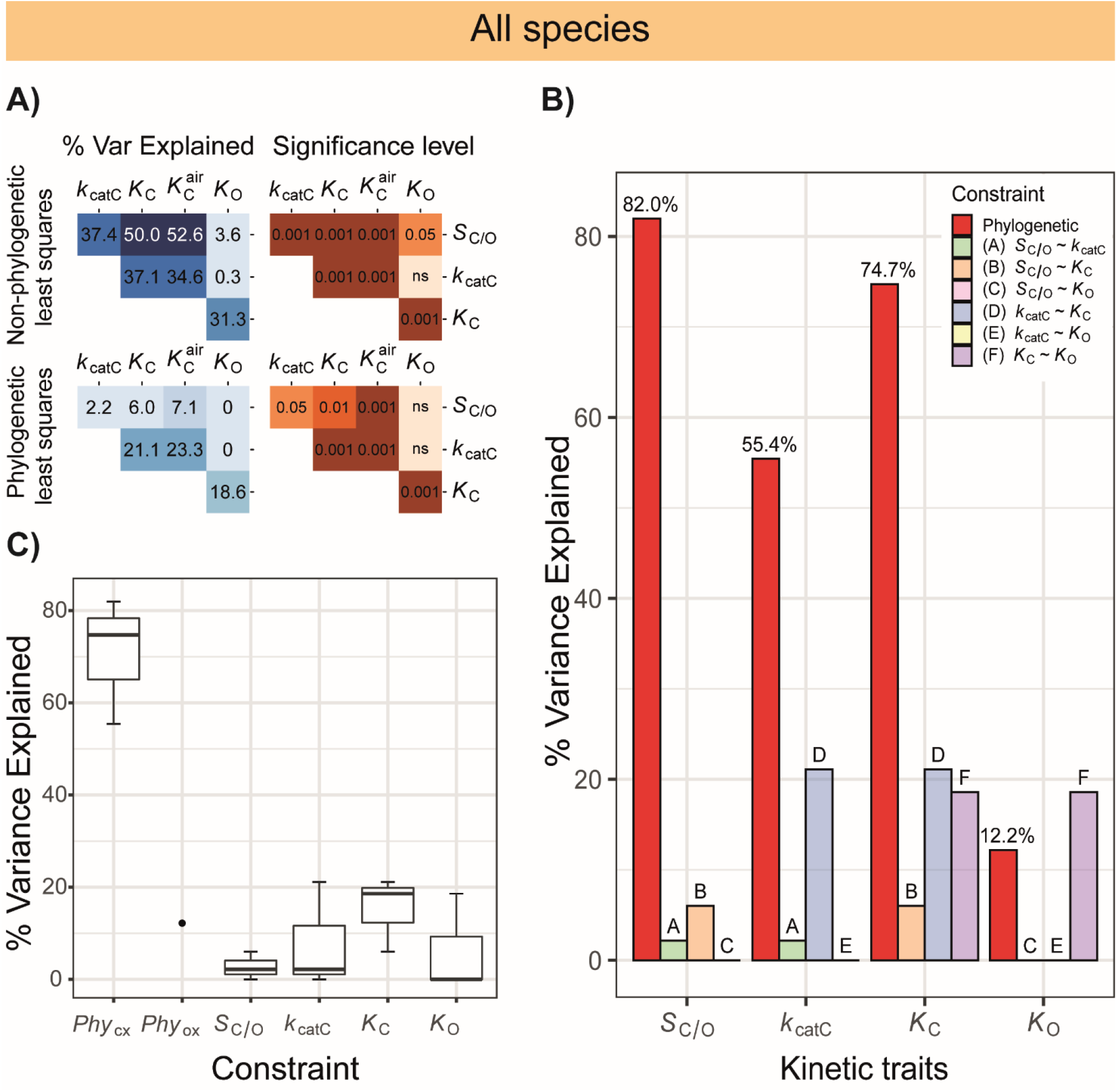
Kinetic and phylogenetic constraints on rubisco adaptation across all studied photosynthetic organisms **A)** Pairwise correlation coefficients (percent variance explained) and associated *p*-values between different rubisco kinetic traits assessed using non-phylogenetic least squares regression models or phylogenetic least squares regression models. It should be noted that the non-corrected correlations measured exhibit slight variation with those reported by (Flamholz *et al*., 2019) owing to the omission of species in this analysis for which *rbcL* sequence data was not available. Significance values are represented as α levels, where; α = 0.001 if *p* < 0.001, α = 0.01 if 0.001 < *p* < 0.01, α = 0.05 if 0.01 < *p* < 0.05, and α = ns if *p* > 0.05. **B)** The variation (%) in rubisco kinetic traits across all photosynthetic organisms that can be explained by phylogenetic constraint and each catalytic trade-off. **C)** Boxplot of all variation explained in each kinetic trait by kinetic trait correlations in comparison to variation explained by phylogeny in all photosynthetic organisms. The phylogenetic constraints on the carboxylase-related traits *Phy*_CX_ (includes *Phy*_*Sc/o*_, *Phy*_*Kcatc*_, and *Phy*_*Kc*_) and phylogenetic constraints on the oxygenase-related trait *Phy*_ox_ (includes *Phy*_*Ko*_ only) are presented separately.

## Discussion

The evolutionary landscape of rubisco has long been proposed to be constrained by catalytic trade-offs. In support of this hypothesis, antagonistic correlations between rubisco kinetic traits inferred from studies comparing limited numbers of species are commonly cited (Tcherkez, Farquhar and Andrews, 2006; Savir *et al*., 2010). Specifically, strong dependencies are thought to occur between rubisco specificity (*S*_C/O_), carboxylase turnover (*k*_catC_) and the Michaelis constants for CO_2_ (*K*_C_) and O_2_ (*K*_O_), respectively (Tcherkez, Farquhar and Andrews, 2006; Savir *et al*., 2010). Combined, these trade-offs are hypothesized to limit the capacity of rubisco to assimilate CO_2_ at high rates by curtailing the inherent carboxylase activity of the enzyme, whilst also causing it to catalyse a reaction with O_2_ which is energetically expensive and results in a loss of fixed carbon (Bowes, Ogren and Hageman, 1971; Chollet, 1977). However, all trade-offs have been inferred based on the assumption that rubisco in different species are independent (Tcherkez, Farquhar and Andrews, 2006; Savir *et al*., 2010; Flamholz *et al*., 2019; Iñiguez *et al*., 2020). Here, we find that this assumption was incorrect and show that a significant phylogenetic signal is found in rubisco kinetic traits across the tree of life. We re-evaluated the extent of rubisco catalytic trade-offs accounting for this phylogenetic signal and found that all catalytic trade-offs were attenuated. The largest trade-offs were observed between the Michaelis constant for CO_2_ and carboxylase turnover (∼21-37 %), and between the Michaelis constants for CO_2_ and O_2_ (∼9-19 %), respectively. Furthermore, we demonstrated that all other catalytic trade-offs were either non-significant or substantially attenuated when the phylogenetic relationship of the species was correctly accounted for. Finally, we found that phylogenetic constraints have played a larger role than catalytic trade-offs in limiting the evolution of rubisco kinetics. Thus, rubisco kinetics have been evolving largely independently of each other in an adaptive landscape that is predominantly limited by phylogenetic constraint.

The presence of a phylogenetic signal in rubisco kinetic traits simply means that rubisco kinetics are more similar among close relatives, with this similarity changing as a function of the phylogenetic distance between species. This result is perhaps not surprising given that all extant rubisco are related by the process of descent with modification from a single ancestral enzyme (Nisbet *et al*., 2007). However, not all biological traits contain a phylogenetic signal (Blomberg, Garland and Ives, 2003; Kamilar and Cooper, 2013). Further, the functional consequences of changes to enzyme sequences are hard to predict (Minshull *et al*., 2005; Damborsky and Brezovsky, 2014; Siegel *et al*., 2015; Carlin *et al*., 2016), with single amino acid substitutions often causing large effects in enzyme kinetics (Cleton-Jansen *et al*., 1991; Villar *et al*., 1997; Johnson *et al*., 2001). Thus, *a priori* it was unknown whether any or all of the rubisco kinetic traits would exhibit a phylogenetic signal. It will be interesting to see whether the presence of phylogenetic signal in enzyme kinetic data is a general phenomenon, and if so it will be important to correctly account for this non-independence when inferring the presence or absence of catalytic trade-offs.

In this work we reveal that the phylogenetic signal in rubisco kinetics is caused by phylogenetic constraint on rubisco that is independent of the catalytic trade-offs. Phylogenetic constraint in this context includes all of the processes that collectively lead to slow rates of molecular evolution. These process include neutral evolution under genetic drift (Felsenstein, 1985), or evolutionary stasis (Prinzing *et al*., 2001; Ackerly, 2004; Cavender-Bares *et al*., 2004; Moles *et al*., 2005; Swenson and Enquist, 2007) under which adaptive change is mitigated by processes including stabilizing selection, pleiotropy and a lack of molecular variability or phenotypic plasticity (Maynard Smith *et al*., 1985; Bradshaw, 1991; Edwards and Naeem, 1993; Wagner, 1995). Although it is possible that multiple factors contribute to the phylogenetic constraint detected in rubisco, it is likely that low molecular variability is a key driver of this phenomenon. For example, the rate of molecular evolution of rubisco is likely constrained by the requirements for i) high levels of transcript and protein abundance (Kelly, 2018; Seward and Kelly, 2018), ii) maintaining complementarity to a wide array of molecular chaperones which assist in protein folding and assembly (e.g., Raf1, Raf2, RbcX, BSD2, Cpn60/Cpn20) and metabolic regulation (e.g., rubisco activase) (Carmo-Silva *et al*., 2015; Aigner *et al*., 2017), and iii) the need to preserve overall protein stability within the molecular activity-stability trade-offs (Studer *et al*., 2014; Duraõ *et al*., 2015; Cummins, Kannappan and Gready, 2018). In angiosperms, these constraints would be further exacerbated due to the presence of the *rbcL* gene in the organellar genome that is uniparentally inherited and does not recombine (Birky, 2001). For example, in angiosperms chloroplast encoded genes evolve 10 times slower than nuclear encoded genes (Wolfe, Li and Sharp, 1987; Smith, 2015). Combined, these evolutionary constraints would hinder the kinetic adaptation of rubisco resulting in the phylogenetic constraint observed in this study.

The strongest catalytic trade-off detected in this study was the ∼21-37 % dependency that was observed between *k*_catC_ and both *K*_C_ and *K*_C_^air^. This finding is compatible with the mechanistic models of rubisco (Farquhar, 1979), and is supported by the recent discovery of enzyme variants which exhibit the highest *k*_catC_ ever recorded at the expense of poor CO_2_ affinities (i.e. *K*_C_ values >250 µM) (Davidi *et al*., 2020). Nevertheless, the dependency between CO_2_ affinity and turnover, despite being the strongest correlation that was observed, is substantially attenuated relative to the coefficients that are conventionally cited (Tcherkez, Farquhar and Andrews, 2006; Savir *et al*., 2010). Therefore, although selecting for a greater rubisco carboxylase turnover is evolutionarily linked with a poorer affinity for CO_2_ (higher *K*_C_), and *vice versa*, significant plasticity exists in this relationship among species such that variation in one kinetic trait only explains ∼21-37 % of variation in the other. This fact explains why there is variability in the carboxylation efficiency among angiosperm rubisco (defined as *k*_catC_/*K*_C_), a core parameter which defines the initial slope of the response of CO_2_ fixation rate to changes in CO_2_ concentration within the aerobic environment of chloroplasts in C_3_ species (Sharwood, 2017). The second strongest catalytic trade-off that was observed was the ∼9-19 % dependency between *K*_C_ and *K*_O_. This trade-off is compatible with the fact that the singular active site of rubisco binds both CO_2_ and O_2_, and thus it is plausible that mutations that affect the active site will affect biding of both substrates, though not necessarily to equivalent extents. All other catalytic trade-offs were either marginal (<9 %) or non-significant. Furthermore, the combined effect of all catalytic trade-offs can only account for 6 - 9 % of total variation in rubisco kinetics between species, a substantially smaller component than can be explained by phylogenetic constraint (30 - 61%).

The phylogenetically resolved analysis of rubisco kinetic evolution also identified changes in kinetic traits associated with the evolution of C_4_ photosynthesis. Specifically, *S*_C/O_ was lower in C_4_ species than in C_3_ species, while *k*_catC_, *K*_C_ and *K*_C_^air^ were higher in C_4_ species than in C_3_ species. Moreover, variation in *K*_O_ was found to be greater in C_4_ species than in C_3_ species. These differences in rubisco kinetics would likely be either neutral or adaptive in a C_4_ context. For example, any change in *K*_O_ would effectively be neutral under the elevated CO_2_ environment of the bundle sheath chloroplast, as it would have only a marginal effect on the *in vivo* carboxylation rate or carboxylation-to-oxygenation ratio, and thus would not cause a concomitant change to organism fitness. In contrast, an increase in *k*_catC_ in the same elevated CO_2_ environment would enable higher flux through rubisco and thus provide an energetic advantage to the cell. Accordingly, one would expect that an increased variation in *K*_O_ in C_4_ species would occur by neutral drift (Kimura, 1991; Savir *et al*., 2010), and that an increased *k*_*cat*C_ would confer a selective advantage even if it came at the expense of a partial reduction in *K*_C_. Thus, the adaptations to rubisco kinetics that occur concomitant with the evolution of C_4_ photosynthesis are consistent with the change in CO_2_:O_2_ ratio and weak catalytic trade-off that exists between *k*_catC_ and *K*_C_. Here, despite the phylogenetic constraints limiting rubisco adaptation, the increased rate at which these kinetic changes occurred in C_4_ species may have been facilitated by the higher rates of molecular evolution (Kelly, 2018) and diversification (Spriggs, Christin and Edwards, 2014) that occurs concomitant with the evolution of C_4_ photosynthesis.

Given the importance of rubisco to life on Earth, the question as to why a “perfect” rubisco has not already evolved is legitimate. For example, although rubisco *K*_C_ is thought to be near optimal in C_3_ plants in light of the ∼8 µM chloroplastic concentration of CO_2_ and the inherent limitations of CO_2_ as a substrate, including its inertness, hydrophobicity and low molecular mass (Andrews and Whitney, 2003; Bar-Even *et al*., 2011; Bathellier *et al*., 2018), the observed *k*_catC_ (∼3 s^−1^ per site) has often been considered low (Bar-Even *et al*., 2011; Tcherkez, 2013; Davidi *et al*., 2018). In addition, all known rubisco variants catalyse a promiscuous and energetically costly reaction with O_2_. However, a recent review of rubisco kinetics relative to those of other enzymes has argued that rubisco is actually not such a bad catalyst (Bathellier *et al*., 2018). Indeed, the phylogenetically informed analysis of rubisco presented here demonstrates that the kinetic traits have been able to evolve largely independently, with kinetic evolution primarily limited by strong phylogenetic constraint. These constraints induce a lag in adaptive evolution that help to explain why the enzyme is better suited to former environmental conditions.

The study presented here highlights the importance of considering phylogenetic relationships when conducing comparative analyses of enzyme kinetics across species. In doing so, it reveals that rubisco evolution has been only weakly constrained by catalytic trade-offs. Instead, phylogenetic constraints, caused by factors that limit the pace of molecular evolution, have provided a more substantial hindrance to rubisco kinetic evolution. Accordingly, it should be feasible in the current synthetic biology revolution to circumvent this evolutionary barrier on rubisco optimisation. Indeed, promising steps toward this goal have been already demonstrated using directed evolution of the enzyme to generate variants with improved catalytic traits in non-photosynthetic archaea (Wilson, Alonso and Whitney, 2016), photosynthetic bacteria (Zhou and Whitney, 2019), and cyanobacteria (Wilson *et al*., 2018). Thus, our findings provide optimism for engineering rubisco in food, fibre and fuel crops to have improved catalytic efficiency.

## Materials and Methods

### Kinetic data

Kinetic measurements of rubisco were attained from (Flamholz *et al*., 2019). *S*_C/O_ values measured using the O_2_ electrode method which calculate [CO_2_] using a pKa of 6.11 (Parry, Keys and Gutteridge, 1989) were normalized relative to *S*_C/O_ values quantified using high precision gas-phase-controlled ^3^H-RuBP-fixation assays (Kane *et al*., 1994) in order to minimise methodological biases in the data. Specifically, as rubisco from wheat (*Triticum aestivum*) was measured in both the O_2_ electrode studies (Orr *et al*., 2016; Prins *et al*., 2016) as well as in the high precision method by (Kane *et al*., 1994), multipliers were applied to all *S*_C/O_ measurements derived from O_2_ electrode assays using wheat as an enzyme standard. The distribution of *S*_C/O_ values in angiosperms before and after normalisation can be seen in (Supplementary File 1, Figure S6).

All kinetic traits in the dataset were transformed on a log scale consistent with (Flamholz *et al*., 2019), and the distributional assumptions of each were verified for analyses herein. For the angiosperm focussed analysis, only angiosperms with experimental measurements of all four principal experimentally measured kinetic traits of interest (*S*_C/O_, *k*_catC_, *K*_C_ and *K*_O_) were taken forward for subsequent analysis. However, all species in the dataset with greater than one kinetic trait measurement were considered for subsequent analyses of all photosynthetic organisms.

Furthermore, a small number of species also had measurements available for *K*_RuBP._ In addition, the measure of the Michaelis constant for CO_2_ under 20.95 % ambient air, *K*_C_^air^, was inferred from *K*_C_ and *K*_O_ based on the formula *K*_C_ + (*K*_C_ [O_2_] / *K*_O_), where 20.95 % [O_2_] in water is 253 µM.

In cases where duplicate entries for a species were present in the kinetic dataset (including synonyms), the median value of their respective kinetic traits was used for inference. In this way, averages were also taken between *Triticum timonovum* and *Triticum timopheevii*, the former of which is a synthetic octoploid of the latter (Murashov and Morozova, 2008). The modified dataset containing corrected *S*_C/O_ values and averages across duplicate entries for species is provided in Supplemental File 2. It should also be noted that values of *k*_*catC*_ are presented as units per site.

### Phylogenetic tree inference

As sequenced genomes or transcriptomes do not exist for many species in the kinetic trait dataset, whole genome phylogenomic approaches could not be used to infer the species tree necessary in order to detect a phylogenetic signal in measured kinetic traits of the rubisco holoenzyme. However, the *rbcL* gene that encodes the large subunit of rubisco has a long history of use for phylogenetic inference of species relationships (Gielly and Taberlet, 1994; APG, 1998, 2016) and was available for all of the angiosperms, and the majority of photosynthetic organisms, that were considered in the analyses. As such, *rbcL* was selected here for use in species tree inference. The coding sequences of *rbcL* for the 137 angiosperm species for which kinetic data was available can be found in Supplementary File 3. The coding sequences of *rbcL* for the complete set of 181 photosynthetic organisms for which both kinetic data and sequencing data was available can be found in Supplementary File 4. Gene sequences were downloaded from NCBI (https://www.ncbi.nlm.nih.gov/) for all species except *Flaveria brownii* which was acquired from the 1KP database (Leebens-Mack *et al*., 2019). Multiple sequence alignments were performed using mafft L-INS-i (Katoh and Standley, 2013), and alignments were trimmed at the terminal ends to remove unaligned positions using AliView (Larsson, 2014). These trimmed nucleotide sequence alignments were used for subsequent phylogenetic analysis. Bootstrapped maximum likelihood phylogenetic trees were inferred by IQ-TREE (Nguyen *et al*., 2015) using the ultrafast bootstrapping method with 1000 replicates and the Shimodaira-Hasegawa approximate likelihood ratio branch test. The best fitting model of sequence evolution was inferred from the data automatically by IQTREE. The resultant maximum likelihood phylogenetic trees were rooted manually using Dendroscope (Huson and Scornavacca, 2012). A number of nodes in the angiosperm tree (Supplementary File 1, Figure S1) and a number of nodes in the tree of all photosynthetic organisms exhibited terminal zero-length branches due to 100 % sequence identify with other closely related species (*n* = 18 and *n* = 23, respectively). These species were condensed into single data points (as their *rbcL* are 100 % identical) and the mean of their kinetic traits was used. This reduced the dataset to 119 unique angiosperm enzymes and 158 unique enzymes across all photosynthetic organisms. The phylogenetic tree inferred from the angiosperm taxa (Supplementary File 1, Figure S1) closely matched the topology of the phylogenetic tree expected from the angiosperm phylogeny with only a few alterations (APG, 2016). Moreover, the topology of the *rbcL* trees most accurately reflects the sequence similarity of the rubisco genes and thus were deemed as suitable for investigation of any phylogenetic signal and its effects on correlations between rubisco kinetic traits.

To confirm that the phylogenetic signal was not attributable to overfitting caused by the use of the *rbcL* gene to infer the phylogeny of rubisco, an analogous maximum likelihood angiosperm phylogenetic tree was inferred using IQ-TREE (Nguyen *et al*., 2015) following the methods described above but based on the angiosperm multiple sequence alignment in which codons containing non-synonymous nucleotide sequence changes were removed (Supplementary File 5). Due to the considerable loss of phylogenetic information accessible for tree building from this alignment, the species tree inferred using nucleotide sequences corresponding to ubiquitously conserved amino acid positions (Supplementary File 1, Figure S2) exhibited an additional number (*n* = 13) of angiosperm species belonging to terminal zero-length branches. As the sequences of these species are known to exhibit non-synonymous mutations which are not included in the tree, it is not appropriate to take means of their kinetic traits as above. As such, these data points were removed from analysis using only this tree. Use of this phylogenetic tree confirmed that the presence of the phylogenetic signal in kinetic traits was not due to overfitting, however as this tree was less accurate than the full-length alignment tree, it was not used for any subsequent analysis.

### Phylogenetic signal analysis

The presence of a phylogenetic signal in kinetic traits was assessed using five different phylogenetic signal detection methods (Gittleman and Kot, 1990; Abouheif, 1999; Pagel, 1999; Blomberg, Garland and Ives, 2003). Here, signal strength was estimated by assessing the distribution of trait values relative to the respective underlying species tree inferred from the *rbcL* sequences using methods which both depend on an explicit evolutionary model, such as Pagel’s lambda (Pagel, 1999) and Blomberg’s K and K* (Blomberg, Garland and Ives, 2003), as well as the spatial autocorrelation analyses of Moran’s I (Gittleman and Kot, 1990) and Abouheif’s Cmean (Abouheif, 1999). Implementation of these phylogenetic signal detection tools was performed using the *phyloSignal* function of the phylosignal package (Keck *et al*., 2016) in the R environment. For further discussion of the differences between phylogenetic signal detection methods see (Münkemüller *et al*., 2012).

### Ancestral state estimation and mapping of kinetic traits to the phylogenetic tree

Ancestral state estimation was conducted to visualise the evolution of rubisco kinetic traits on the phylogenetic tree which relates the angiosperms. For this purpose, the kinetic dataset was mapped and scaled onto the angiosperm species tree by employing the function *ggtree* in the ggtree package (Yu *et al*., 2017). Here, terminal branches were coloured according to the measurement of the kinetic trait in the species which comprise the terminal branch, whereas internal branches were coloured based on values inferred in ancestral species using ancestral state estimation (Yu *et al*., 2017).

### Least squares and linear regression models

All regression models between pairwise combinations of kinetic traits were fit in the R environment. Phylogenetic generalized regression accounting for the phylogenetic non-independence between species was performed using the function *pgls* in the caper package (Comparative Analyses of Phylogenetics and Evolution in R) (Orme *et al*., 2014). In each case, the structure of the phylogenetic signal was corrected for using branch length transformations of the phylogenetic tree based on the mean maximum likelihood estimates of lambda calculated for the traits being interrogated, with kapa and delta held constant. In cases where the mean maximum likelihood estimates of lambda exceeded the upper limit of the model, this value was set to 1. Phylogenetic corrections to differences in kinetic trait values between C_3_ and C_4_ plants based on the phylogenetic non-independence of species were also applied using the *pgls* function in the caper package (Orme *et al*., 2014) with photosynthetic type incorporated as a factorial variable.

In order to partition the variance in rubisco kinetic traits explained by phylogenetic constraints as compared to non-phylogenetic constraints, the rr2 package (Ives and Li, 2018) was employed in R. Here, to assess the contribution of phylogeny in explaining the variation in kinetic trait values, the explanatory power of the phylogenetic component was measured by comparing full and reduced phylogenetic regression models using the partial *R*^2^_pred_ inferential statistical, based on advice from (Ives, 2019) of this being the most intuitive and direct approach to understand how different model components explain variation. For this analysis, phylogenetic regression models were fit using the *phylolm* function in the phylolm package (Tung Ho and Ané, 2014) using Pagel’s lambda model for the error term.

## Acknowledgements

The authors would like to thank Avi Flamholz for his comments on the manuscript. This work was funded by the Royal Society and the European Union’s Horizon 2020 research and innovation program under grant agreement number 637765. JWB, JRN, JSB, AE, AB and AU were funded by the Biotechnology and Biological Sciences Research Council (BBSRC) [grant numbers BB/M011224/1 and BB/P003117/1]. TR and SMW were supported by the Australian Government through the Australian Research Council Centre of Excellence for Translational Photosynthesis CE140100015.

## Data Availability

All data used in this study is provided in the supplemental material.

## Author Contributions

SK conceived the study. JWB conducted the analysis. JWB, TR, SMW, DME, and SK analyzed the data. SK and JWB wrote the manuscript. JRN, JSB, AE, AB and AU contributed to discussions and methodological development. All authors edited the manuscript.

## References

Abouheif, E. (1999) A method for testing the assumption of phylogenetic independence in comparative data, Evolutionary Ecology Research. doi: 10.1007/s11295-009-0238-5.

Ackerly, D. D. (2004) ‘Adaptation, niche conservatism, and convergence: Comparative studies of leaf evolution in the California chaparral’, American Naturalist, 163(5), pp. 654–671. doi: 10.1086/383062.

Aigner, H. et al. (2017) ‘Plant RuBisCo assembly in E. coli with five chloroplast chaperones including BSD2’, Science. doi: 10.1126/science.aap9221.

Andersson, I. and Backlund, A. (2008) ‘Structure and function of Rubisco’, Plant Physiology and Biochemistry, 46(3), pp. 275–291. doi: 10.1016/j.plaphy.2008.01.001.

Andrews, T. J. (1988) ‘Catalysis by cyanobacterial ribulose-bisphosphate carboxylase large subunits in the complete absence of small subunits.’, The Journal of biological chemistry.

Andrews, T. J. and Whitney, S. M. (2003) ‘Manipulating ribulose bisphosphate carboxylase/oxygenase in the chloroplasts of higher plants’, Archives of Biochemistry and Biophysics, 414(2), pp. 159–169. doi: 10.1016/S0003-9861(03)00100-0.

APG (1998) An Ordinal Classification for the Families of Flowering Plants, Annals of the Missouri Botanical Garden. doi: 10.2307/2992015.

APG (2016) ‘An update of the Angiosperm Phylogeny Group classification for the orders and families of flowering plants: APG IV’, Botanical Journal of the Linnean Society. Oxford University Press, 181(1), pp. 1–20. doi: 10.1111/boj.12385.

Araújo, W. L. et al. (2017) ‘Engineering photosynthesis: Progress and perspectives’, F1000Research. doi: 10.12688/f1000research.12181.1.

Badger, M. R. and Lorimer, G. H. (1976) ‘Activation of ribulose-1,5-bisphosphate oxygenase. The role of Mg2+, CO2, and pH’, Archives of Biochemistry and Biophysics, pp. 723–729. doi: 10.1016/0003-9861(76)90565-8.

Banda, D. M. et al. (2020) ‘Novel bacterial clade reveals origin of form I Rubisco’, Nature Plants. doi: 10.1038/s41477-020-00762-4.

Bar-Even, A. et al. (2011) ‘The moderately efficient enzyme: Evolutionary and physicochemical trends shaping enzyme parameters’, Biochemistry, 50(21), pp. 4402–4410. doi: 10.1021/bi2002289.

Bar-On, Y. M. and Milo, R. (2019) ‘The global mass and average rate of rubisco’, Proceedings of the National Academy of Sciences of the United States of America. PNAS, 116(10), pp. 4738–4743. doi: 10.1073/pnas.1816654116.

Bathellier, C. et al. (2018) ‘Rubisco is not really so bad’, Plant Cell and Environment, pp. 705–716. doi: 10.1111/pce.13149.

Bauwe, H. et al. (2012) ‘Photorespiration has a dual origin and manifold links to central metabolism’, Current Opinion in Plant Biology, pp. 269–275. doi: 10.1016/j.pbi.2012.01.008.

Beer, C. et al. (2010) ‘Terrestrial gross carbon dioxide uptake: Global distribution and covariation with climate’, Science, 329(5993), pp. 834–838. doi: 10.1126/science.1184984.

Birky, C. W. (2001) ‘The Inheritance of Genes in Mitochondria and Chloroplasts: Laws, Mechanisms, and Models’, Annual Review of Genetics. doi: 10.1146/annurev.genet.35.102401.090231.

Blomberg, S. P., Garland, T. and Ives, A. R. (2003) ‘Testing for phylogenetic signal in comparative data: Behavioral traits are more labile’, Evolution. Oxford, UK: Blackwell Publishing Ltd, pp. 717– 745. doi: 10.1111/j.0014-3820.2003.tb00285.x.

Bowes, G., Ogren, W. L. and Hageman, R. H. (1971) Phosphoglycolate production catalyzed by ribulose diphosphate carboxylase, Biochemical and Biophysical Research Communications. doi: 10.1016/0006-291X(71)90475-X.

Bradshaw, A. (1991) ‘Genostasis and the limits to evolution’, Philosophical transactions of the Royal Society of London. Series B, Biological sciences, 333(1267), pp. 289–305.

Busch, F. A. (2020) ‘Photorespiration in the context of Rubisco biochemistry, CO2 diffusion and metabolism’, Plant Journal, 101(4), pp. 919–939. doi: 10.1111/tpj.14674.

Carlin, D. A. et al. (2016) ‘Kinetic characterization of 100 glycoside hydrolase mutants enables the discovery of structural features correlated with kinetic constants’, PLoS ONE. doi: 10.1371/journal.pone.0147596.

Carmo-Silva, E. et al. (2015) ‘Optimizing Rubisco and its regulation for greater resource use efficiency’, Plant, Cell and Environment, pp. 1817–1832. doi: 10.1111/pce.12425.

Cavender-Bares, J. et al. (2004) ‘Phylogenetic overdispersion in Floridian oak communities’, American Naturalist, 163(6), pp. 823–843. doi: 10.1086/386375.

Cheverud, J. M., Dow, M. M. and Leutenegger, W. (1985) ‘the Quantitative Assessment of Phylogenetic Constraints in Comparative Analyses: Sexual Dimorphism in Body Weight Among Primates’, Evolution. Wiley, 39(6), pp. 1335–1351. doi: 10.1111/j.1558-5646.1985.tb05699.x.

Cheverud, J. M., Dow, M. M. and Leutenegger, W. (1986) ‘A Phylogenetic Autocorrelation Analysis of Sexual Dimorphism in Primates’, American Anthropologist. Wiley-Blackwell, 88(4), pp. 916–922. doi: 10.1525/aa.1986.88.4.02a00090.

Chollet, R. (1977) The biochemistry of photorespiration, Trends in Biochemical Sciences. doi: 10.1016/0968-0004(77)90364-4.

Chollet, R. and Anderson, L. L. (1976) ‘Regulation of ribulose 1,5-bisphosphate carboxylase-oxygenase activities by temperature pretreatment and chloroplast metabolites’, Archives of Biochemistry and Biophysics, pp. 344–351. doi: 10.1016/0003-9861(76)90173-9.

Cleton-Jansen, A. M. et al. (1991) ‘A single amino acid substitution changes the substrate specificity of quinoprotein glucose dehydrogenase in Gluconobacter oxydans’, MGG Molecular & General Genetics. doi: 10.1007/BF00272157.

Cummins, P. L., Kannappan, B. and Gready, J. E. (2018) ‘Directions for optimization of photosynthetic carbon fixation: Rubisco’s efficiency may not be so constrained after all’, Frontiers in Plant Science, 9. doi: 10.3389/fpls.2018.00183.

Damborsky, J. and Brezovsky, J. (2014) ‘Computational tools for designing and engineering enzymes’, Current Opinion in Chemical Biology. doi: 10.1016/j.cbpa.2013.12.003.

Davidi, D. et al. (2018) ‘A Bird’s-Eye View of Enzyme Evolution: Chemical, Physicochemical, and Physiological Considerations’, Chemical Reviews, 118(18), pp. 8786–8797. doi: 10.1021/acs.chemrev.8b00039.

Davidi, D. et al. (2020) ‘Highly active rubiscos discovered by systematic interrogation of natural sequence diversity’, The EMBO Journal. John Wiley & Sons, Ltd. doi: 10.15252/embj.2019104081.

Duraõ, P. et al. (2015) ‘Opposing effects of folding and assembly chaperones on evolvability of Rubisco’, Nature Chemical Biology, 11(2), pp. 148–155. doi: 10.1038/nchembio.1715.

Eckardt, N. A. (2005) ‘Photorespiration revisited’, Plant Cell. American Society of Plant Biologists, pp. 2139–2141. doi: 10.1105/tpc.105.035873.

Edwards, S. V. and Naeem, S. (1993) ‘The phylogenetic component of cooperative breeding in perching birds’, American Naturalist. Am Nat, 141(5), pp. 754–789. doi: 10.1086/285504.

Ehleringer, J. R. et al. (1991) Climate change and the evolution of C4 photosynthesis, Trends in Ecology and Evolution. doi: 10.1016/0169-5347(91)90183-X.

Ellis, R. J. (1979) The most abundant protein in the world, Trends in Biochemical Sciences. doi: 10.1016/0968-0004(79)90212-3.

Erb, T. J. and Zarzycki, J. (2018) A short history of RubisCO: the rise and fall (?) of Nature’s predominant CO2 fixing enzyme, Current Opinion in Biotechnology. doi: 10.1016/j.copbio.2017.07.017.

Farquhar, G. D. (1979) ‘Models describing the kinetics of ribulose biphosphate carboxylase-oxygenase’, Archives of Biochemistry and Biophysics, 193(2), pp. 456–468. doi: 10.1016/0003-9861(79)90052-3.

Felsenstein, J. (1985) Phylogenies and the comparative method., American Naturalist. doi: 10.1086/284325.

Flamholz, A. I. et al. (2019) ‘Revisiting Trade-offs between Rubisco Kinetic Parameters’, Biochemistry, pp. 3365–3376. doi: 10.1021/acs.biochem.9b00237.

Flamholz, A. and Shih, P. M. (2020) ‘Cell biology of photosynthesis over geologic time’, Current Biology. doi: 10.1016/j.cub.2020.01.076.

Fukayama, H. et al. (2019) ‘Rubisco small subunits of C4 plants, Napier grass and guinea grass confer C4-like catalytic properties on Rubisco in rice’, Plant Production Science. doi: 10.1080/1343943X.2018.1540279.

Garland, T. (2001) ‘Phylogenetic comparison and artificial selection’, in. Springer, Boston, MA, pp. 107–132. doi: 10.1007/978-1-4757-3401-0_9.

Gielly, L. and Taberlet, P. (1994) The use of chloroplast DNA to resolve plant phylogenies: noncoding versus rbcL sequences., Molecular Biology and Evolution. doi: 10.1093/oxfordjournals.molbev.a040157.

Gittleman, J. L. and Kot, M. (1990) ‘Adaptation: Statistics and a Null Model for Estimating Phylogenetic Effects’, Systematic Zoology. Society of Systematic Zoology, pp. 227–241. doi: 10.2307/2992183.

Grafen, A. (1989) ‘The phylogenetic regression.’, Philosophical transactions of the Royal Society of London. Series B, Biological sciences. doi: 10.1098/rstb.1989.0106.

Gunn, L. H. et al. (2020) ‘The dependency of red Rubisco on its cognate activase for enhancing plant photosynthesis and growth’, Proceedings of the National Academy of Sciences of the United States of America. doi: 10.1073/pnas.2011641117.

Hardy, O. J. and Pavoine, S. (2012) ‘Assessing phylogenetic signal with measurement error: A comparison of mantel tests, blomberg et al.’s K, and phylogenetic distograms’, Evolution. John Wiley & Sons, Ltd, 66(8), pp. 2614–2621. doi: 10.1111/j.1558-5646.2012.01623.x.

Huson, D. H. and Scornavacca, C. (2012) ‘Dendroscope 3: An interactive tool for rooted phylogenetic trees and networks’, Systematic Biology. Oxford University Press, pp. 1061–1067. doi: 10.1093/sysbio/sys062.

Iñiguez, C. et al. (2020) ‘Evolutionary trends in RuBisCO kinetics and their co-evolution with CO2 concentrating mechanisms’, Plant Journal, 101(4), pp. 897–918. doi: 10.1111/tpj.14643.

Ishikawa, C. et al. (2011) ‘Functional incorporation of sorghum small subunit increases the catalytic turnover rate of rubisco in transgenic rice’, Plant Physiology. doi: 10.1104/pp.111.177030.

Ives, A. and Li, D. (2018) ‘rr2: An R package to calculate $R^2$s for regression models’, Journal of Open Source Software, 3(30), p. 1028. doi: 10.21105/joss.01028.

Ives, A. R. (2019) ‘R 2 s for Correlated Data: Phylogenetic Models, LMMs, and GLMMs’, Systematic Biology, 68(2), pp. 234–251. doi: 10.1093/sysbio/syy060.

Johnson, E. T. et al. (2001) ‘Alteration of a single amino acid changes the substrate specificity of dihydroflavonol 4-reductase’, Plant Journal. doi: 10.1046/j.1365-313X.2001.00962.x.

Joshi, J. et al. (2015) ‘Role of small subunit in mediating assembly of red-type Form I Rubisco’, Journal of Biological Chemistry. doi: 10.1074/jbc.M114.613091.

Kamilar J. M. and Cooper, N. (2013) ‘Phylogenetic signal in primate behaviour, ecology and life history’, Philosophical Transactions of the Royal Society B: Biological Sciences, 368(1618). doi: 10.1098/rstb.2012.0341.

Kane, H. J. et al. (1994) ‘An improved method for measuring the CO2/O2 specificity of ribulosebisphosphatte carboxylase-oxygenase’, Australian Journal of Plant Physiology. doi: 10.1071/PP9940449.

Katoh, K. and Standley, D. M. (2013) ‘MAFFT multiple sequence alignment software version 7: Improvements in performance and usability’, Molecular Biology and Evolution. Oxford University Press, pp. 772–780. doi: 10.1093/molbev/mst010.

Keck, F. et al. (2016) ‘Phylosignal: An R package to measure, test, and explore the phylogenetic signal’, Ecology and Evolution, pp. 2774–2780. doi: 10.1002/ece3.2051.

Kelly, S. (2018) ‘The amount of nitrogen used for photosynthesis modulates molecular evolution in plants’, Molecular Biology and Evolution, 35(7), pp. 1616–1625. doi: 10.1093/molbev/msy043.

Kimura, M. (1991) ‘The neutral theory of molecular evolution: A review of recent evidence’, The Japanese Journal of Genetics. doi: 10.1266/jjg.66.367.

Larsson, A. (2014) ‘AliView: A fast and lightweight alignment viewer and editor for large datasets’, Bioinformatics. Oxford University Press, pp. 3276–3278. doi: 10.1093/bioinformatics/btu531.

Lee, B., Berka, R. M. and Tabita F. R. (1991) ‘Mutations in the small subunit of cyanobacterial ribulose-bisphosphate carboxylase/oxygenase that modulate interactions with large subunits’, Journal of Biological Chemistry.

Lee, B. and Tabita, F. R. (1990) ‘Purification of Recombinant Ribulose-l,5-bisphosphate Carboxylase/Oxygenase Large Subunits Suitable for Reconstitution and Assembly of Active L8S8 Enzyme’, Biochemistry. doi: 10.1021/bi00492a007.

Leebens-Mack, J. H. et al. (2019) ‘One thousand plant transcriptomes and the phylogenomics of green plants’, Nature. doi: 10.1038/s41586-019-1693-2.

Martin-Avila, E. et al. (2020) ‘Modifying Plant Photosynthesis and Growth via Simultaneous Chloroplast Transformation of Rubisco Large and Small Subunits.’, The Plant cell. doi: 10.1105/tpc.20.00288.

Maynard Smith, J. et al. (1985) ‘Developmental constraints on evolution’, The Quarterly Review of Biology.

Meyer, M. and Griffiths, H. (2013) ‘Origins and diversity of eukaryotic CO2-concentrating mechanisms: Lessons for the future’, Journal of Experimental Botany, pp. 769–786. doi: 10.1093/jxb/ers390.

Minshull, J. et al. (2005) ‘Predicting enzyme function from protein sequence’, Current Opinion in Chemical Biology. doi: 10.1016/j.cbpa.2005.02.003.

Moles, A. T. et al. (2005) ‘A brief history of seed size’, Science, 307(5709), pp. 576–580. doi: 10.1126/science.1104863.

Münkemüller, T. et al. (2012) ‘How to measure and test phylogenetic signal’, Methods in Ecology and Evolution, 3(4), pp. 743–756. doi: 10.1111/j.2041-210X.2012.00196.x.

Murashov, V. V. and Morozova Z. A. (2008) ‘Comparative morphogenesis of Triticum timopheevii (Zhuk.) and synthetic octoploid species T. timonovum Heslot et Ferrary’, Moscow University Biological Sciences Bulletin. Heidelberg: Allerton Press, Inc., pp. 127–133. doi: 10.3103/s0096392508030073.

Nevo, O. et al. (2020) ‘The evolution of fruit scent: phylogenetic and developmental constraints’, BMC Evolutionary Biology. doi: 10.1186/s12862-020-01708-2.

Nguyen, L. T. et al. (2015) ‘IQ-TREE: A fast and effective stochastic algorithm for estimating maximum-likelihood phylogenies’, Molecular Biology and Evolution, pp. 268–274. doi: 10.1093/molbev/msu300.

Nisbet, E. G. et al. (2007) ‘The age of Rubisco: The evolution of oxygenic photosynthesis’, Geobiology, 5(4), pp. 311–335. doi: 10.1111/j.1472-4669.2007.00127.x.

Ogren, W. L. (1984) ‘Photorespiration: Pathways, Regulation, and Modification’, Annual Review of Plant Physiology. Annual Reviews, 35(1), pp. 415–442. doi: 10.1146/annurev.pp.35.060184.002215.

Ogren, W. L. and Bowes, G. (1971) ‘Ribulose diphosphate carboxylase regulates soybean photorespiration’, Nature New Biology. Nat New Biol, 230(13), pp. 159–160. doi: 10.1038/newbio230159a0.

Orme, D. et al. (2014) Caper: Comparative analyses of phylogenetics and evolution in R, R package version 0.5.2/ r121. Available at: http://r-forge.r-project.org/projects/caper/ (Accessed: 31 May 2020).

Orr, D. J. et al. (2016) ‘Surveying rubisco diversity and temperature response to improve crop photosynthetic efficiency’, Plant Physiology. doi: 10.1104/pp.16.00750.

Pagel, M. (1999) ‘Inferring the historical patterns of biological evolution’, Nature. Nature Publishing Group, pp. 877–884. doi: 10.1038/44766.

Pagel M. D. and Harvey P. H. (1989) ‘Comparative methods for examining adaptation depend on evolutionary models.’, Folia primatologica; international journal of primatology. doi: 10.1159/000156417.

Parry, M. A. et al. (2013) ‘Rubisco activity and regulation as targets for crop improvement | Journal of Experimental Botany | Oxford Academic’, Journal of Experimental Bitany.

Parry, M. A. J. et al. (2007) ‘Prospects for increasing photosynthesis by overcoming the limitations of Rubisco’, in Journal of Agricultural Science. doi: 10.1017/S0021859606006666.

Parry, M. A. J., Keys, A. J. and Gutteridge, S. (1989) ‘Variation in the specificity factor of C3 higher plant rubiscos determined by the total consumption of ribulose-P2’, Journal of Experimental Botany. doi: 10.1093/jxb/40.3.317.

Peterhansel, C. et al. (2010) ‘Photorespiration’, The Arabidopsis Book. BioOne, 8(8), p. e0130. doi: 10.1199/tab.0130.

Prins, A. et al. (2016) ‘Rubisco catalytic properties of wild and domesticated relatives provide scope for improving wheat photosynthesis’, Journal of Experimental Botany. doi: 10.1093/jxb/erv574.

Prinzing, A. et al. (2001) ‘The niche of higher plants: Evidence for phylogenetic conservatism’, Proceedings of the Royal Society B: Biological Sciences, 268(1483), pp. 2383–2389. doi: 10.1098/rspb.2001.1801.

Read, B. A. and Tabita, F. R. (1992a) ‘A Hybrid Ribulosebisphosphate Carboxylase/Oxygenase Enzyme Exhibiting a Substantial Increase in Substrate Specificity Factor’, Biochemistry. doi: 10.1021/bi00139a018.

Read B. A. and Tabita F. R. (1992b) ‘Amino Acid Substitutions in the Small Subunit of Ribulose-1, 5-bisphosphate Carboxylase/Oxygenase That Influence Catalytic Activity of the Holoenzyme’, Biochemistry. doi: 10.1021/bi00117a031.

Savir, Y. et al. (2010) ‘Cross-species analysis traces adaptation of Rubisco toward optimality in a low-dimensional landscape’, Proceedings of the National Academy of Sciences of the United States of America, 107(8), pp. 3475–3480. doi: 10.1073/pnas.0911663107.

Schneider, G., Lindqvist, Y. and Brändén, C. I. (1992) ‘RUBISCO: Structure and mechanism’, Annual Review of Biophysics and Biomolecular Structure. doi: 10.1146/annurev.bb.21.060192.001003.

Seward, E. A. and Kelly, S. (2018) ‘Selection-driven cost-efficiency optimization of transcripts modulates gene evolutionary rate in bacteria’, Genome Biology, 19(1). doi: 10.1186/s13059-018-1480-7.

Sharkey, T. D. (2020) ‘Emerging research in plant photosynthesis’, Emerging Topics in Life Sciences. Portland Press Ltd. doi: 10.1042/etls20200035.

Sharwood, R. E. (2017) ‘Engineering chloroplasts to improve Rubisco catalysis: prospects for translating improvements into food and fiber crops’, New Phytologist, pp. 494–510. doi: 10.1111/nph.14351.

Sharwood, R. E., Ghannoum, O. and Whitney, S. M. (2016) ‘Prospects for improving CO2 fixation in C3-crops through understanding C4-Rubisco biogenesis and catalytic diversity’, Current Opinion in Plant Biology. doi: 10.1016/j.pbi.2016.04.002.

Siegel, J. B. et al. (2015) ‘Computational protein design enables a novel one-carbon assimilation pathway’, Proceedings of the National Academy of Sciences of the United States of America. doi: 10.1073/pnas.1500545112.

Smith, D. R. (2015) ‘Mutation rates in plastid genomes: They are lower than you might think’, Genome Biology and Evolution. doi: 10.1093/gbe/evv069.

Spreitzer, R. J., Peddi, S. R. and Satagopan, S. (2005) ‘Phylogenetic engineering at an interface between large and small subunits imparts land-plant kinetic properties to algal Rubisco’ Proceedings of the National Academy of Sciences of the United States of America. doi: 10.1073/pnas.0508042102.

Spriggs, E. L., Christin, P. A. and Edwards, E. J. (2014) ‘C4 photosynthesis promoted species diversification during the miocene grassland expansion’, PLoS ONE. doi: 10.1371/journal.pone.0097722.

Studer, R. A. et al. (2014) ‘Stability-activity tradeoffs constrain the adaptive evolution of RubisCO’, Proceedings of the National Academy of Sciences of the United States of America, 111(6), pp. 2223– 2228. doi: 10.1073/pnas.1310811111.

Su, Z. and Townsend, J. P. (2015) ‘Utility of characters evolving at diverse rates of evolution to resolve quartet trees with unequal branch lengths: Analytical predictions of long-branch effects’, BMC Evolutionary Biology, p. 86. doi: 10.1186/s12862-015-0364-7.

Swenson, N. G. and Enquist, B. J. (2007) ‘Ecological and evolutionary determinants of a key plant functional trait: Wood density and its community-wide variation across latitude and elevation’, American Journal of Botany, 94(3), pp. 451–459. doi: 10.3732/ajb.94.3.451.

Tabita, F. R. et al. (2008) ‘Distinct form I, II, III, and IV Rubisco proteins from the three kingdoms of life provide clues about Rubisco evolution and structure/function relationships’, Journal of Experimental Botany, 59(7), pp. 1515–1524. doi: 10.1093/jxb/erm361.

Tcherkez, G. (2013) ‘Modelling the reaction mechanism of ribulose-1,5-bisphosphate carboxylase/oxygenase and consequences for kinetic parameters’, Plant, Cell and Environment, 36(9), pp. 1586–1596. doi: 10.1111/pce.12066.

Tcherkez, G. G. B., Farquhar, G. D. and Andrews, T. J. (2006) ‘Despite slow catalysis and confused substrate specificity, all ribulose bisphosphate carboxylases may be nearly perfectly optimized’, Proceedings of the National Academy of Sciences of the United States of America, 103(19), pp. 7246–7251. doi: 10.1073/pnas.0600605103.

Tung Ho L. S. and Ané, C. (2014) ‘A linear-time algorithm for gaussian and non-gaussian trait evolution models’, Systematic Biology, 63(3), pp. 397–408. doi: 10.1093/sysbio/syu005.

Villar K. Del et al. (1997) ‘Amino acid substitutions that convert the protein substrate specificity of farnesyltransferase to that of geranylgeranyltransferase type I’, Journal of Biological Chemistry. doi: 10.1074/jbc.272.1.680.

Wagner, P. J. (1995) ‘Testing Evolutionary Constraint Hypotheses with Early Paleozoic Gastropods’, Paleobiology. doi: 10.1017/S0094837300013294.

Whitney, S. M., Houtz, R. L. and Alonso, H. (2011) ‘Advancing our understanding and capacity to engineer nature’s CO2-sequestering enzyme, Rubisco’, Plant Physiology, 155(1), pp. 27–35. doi: 10.1104/pp.110.164814.

Wilson, R. H. et al. (2018) ‘An improved Escherichia coli screen for Rubisco identifies a protein-protein interface that can enhance CO2-fixation kinetics’, Journal of Biological Chemistry. doi: 10.1074/jbc.M117.810861.

Wilson, R. H., Alonso, H. and Whitney, S. M. (2016) ‘Evolving Methanococcoides burtonii archaeal Rubisco for improved photosynthesis and plant growth’, Scientific Reports. doi: 10.1038/srep22284.

Wolfe, K. H., Li, W. H. and Sharp, P. M. (1987) ‘Rates of nucleotide substitution vary greatly among plant mitochondrial, chloroplast, and nuclear DNAs.’, Proceedings of the National Academy of Sciences of the United States of America. doi: 10.1073/pnas.84.24.9054.

Yu, G. et al. (2017) ‘ggtree: an r package for visualization and annotation of phylogenetic trees with their covariates and other associated data’, Methods in Ecology and Evolution. doi: 10.1111/2041-210X.12628.

Zhou, Y. and Whitney, S. (2019) ‘Directed evolution of an improved Rubisco; in vitro analyses to decipher fact from fiction’, International Journal of Molecular Sciences, 20(20). doi: 10.3390/ijms20205019.

