## Supplemental File S1 for "Rubisco adaptation is more limited by phylogenetic constraint than by catalytic trade-off"

### Supplementary File 1

#### The phylogenetic signal in rubisco kinetic traits is not caused by overfitting due to use of rbcL for phylogenetic tree inference

Although a phylogenetic tree inferred from the nucleotide sequence of the *rbcL* gene is widely accepted as a good proxy for the true phylogenetic relationships between plant species (Gielly and Taberlet, 1994), and is used in many large scale studies of plant phylogeny (APG, 2016), there is a possibility that the use of this gene when assessing for a phylogenetic signal in rubisco kinetic traits could have resulted in overfitting. Specifically, amino acid substitutions that contribute to variation in measured rubisco kinetics would also contribute to the topology of the phylogenetic tree. Thus, to test for the presence of this potential overfitting and prevent such amino acid substitutions influencing the phylogenetic relationship between species, a phylogenetic tree was inferred for all species from a nucleotide sequence alignment of the *rbcL* gene using only ubiquitously conserved amino acid positions (i.e. sites where the amino acid was constant across all species, but the coding sequence exhibited synonymous sequence changes). The use of this phylogenetic tree containing no amino acid sequence variation (Figure S2 below) did not affect the conclusions of phylogenetic signal methods in any kinetic trait assessed. Here, an overall significant phylogenetic signal was found in all kinetic traits except *K*_O_ (Table S1 below), analogous to results previously described based on the tree inferred from the complete *rbcL* coding sequence (Table 1). Therefore, measurements of the phylogenetic signal are not an artefact of using the *rbcL* gene to infer the phylogenetic tree.

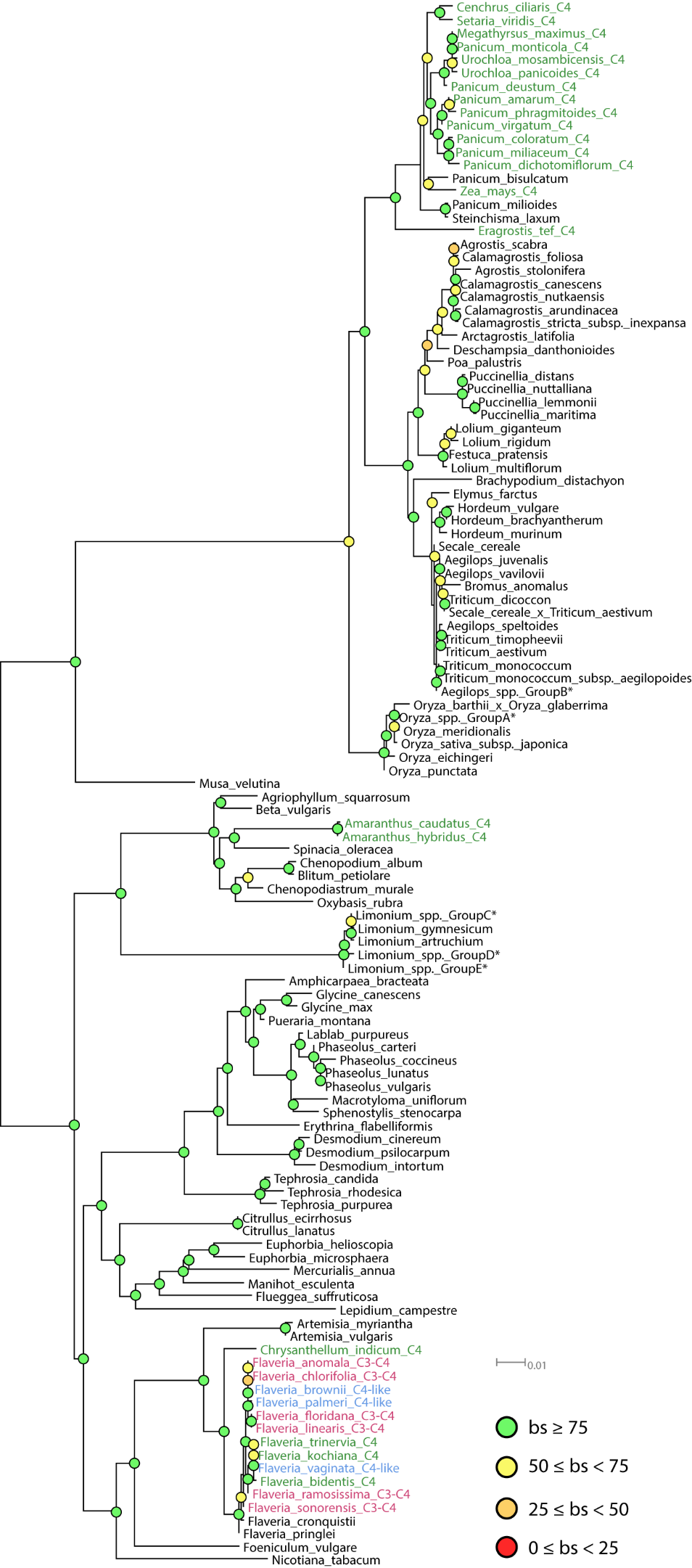

**Supplementary Figure S1.**  Consensus phylogenetic tree of the angiosperm species in this study inferred from the nucleotide sequence of the *rbcL* gene of the large subunit of the rubisco enzyme. Bootstrap support (bs) values are indicated at internal nodes with scale indicated next to the tree. Where known, photosynthetic types other than the ancestral C_3_ pathway have been annotated on to species labels which are colour coded for visualisation, and include C_3_-C_4_ intermediates (red), C_4_-like (light green) and C_4_ plants (green). Species which shared identical gene sequences have been condensed into single nodes on the tree to avoid terminal zero-length branches and are signified by asterisks. Of these, *Oryza* spp. Group A include *O. longistaminata*, *O. glaberrima*, *O. sativa* f. *spontanea*, *O. sativa* subpsp. *indica* and *O. glumipatula*; *Aegilops* spp. Group B include *A. triuncialis*, *A. uniaristata*, *A. tauschii*, *A. comosa*, *A.* *biuncialis*, *A. cylindrica*; *Limonium* spp. Group C include *L. antonii-llorensii*, *L. gibertii*, *L. biflorum*; *Limonium* spp. Group D include *L.* *echioides*, *L. barceloi*, *L.* *balearicum*, *L. companyonis*; and *Limonium* spp. Group E include *L. ejulabilis*, *L. retusum*, *L. leonardi-llorensii*, *L. magallufianum*, *L. grosii*.

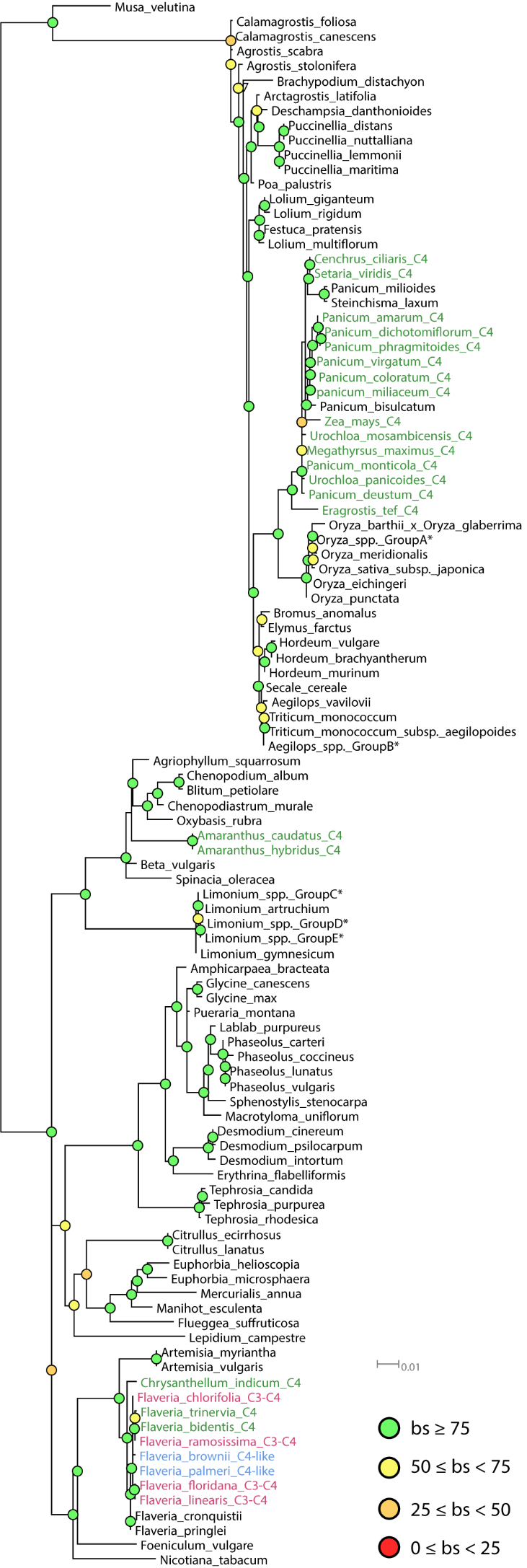

**Supplementary Figure S2.** Consensus phylogenetic tree of the angiosperm species in this study inferred from the nucleotide sequence of the *rbcL* gene of the large subunit of the rubisco enzyme when including only sites which encode ubiquitously conserved amino acid positions. The confidence of branch points on the tree are depicted as nodes which are colour coded by bootstrap support (bs) values. Where known, photosynthetic types other than the ancestral C_3_ pathway have been annotated on to species labels which are colour coded for visualisation, and include C_3_-C_4_ intermediates (red), C_4_-like (blue) and C_4_ plants (green). Species which shared identical sequences when considering the entire sequence length have been condensed into single nodes on the tree to avoid terminal zero-length branches and are signified by asterisks (see supplementary Figure S1 for further details). In addition to these, a number of terminal zero length branches were present between nodes on the tree as an artefact of removing non-synonymous positions in the *rbcL* sequence alignment. All but one of the species within these respective groups of plants containing zero length branches were removed and are thus not present in this tree, including *Aegilops juvenalis*, *Aegilops speltoides*, *Calamagrostis arundinacea*, *Calamagrostis stricta subsp. inexpansa*, *Calamagrostis nutkaensis*, *Flaveria anomala*, *Flaveria kochiana*, *Flaveria sonorensis*, *Flaveria vaginata*, *Triticum dicoccon*, *Triticum timopheevii*, *Secale cereale x Triticum aestivum* and *Triticum aestivum*

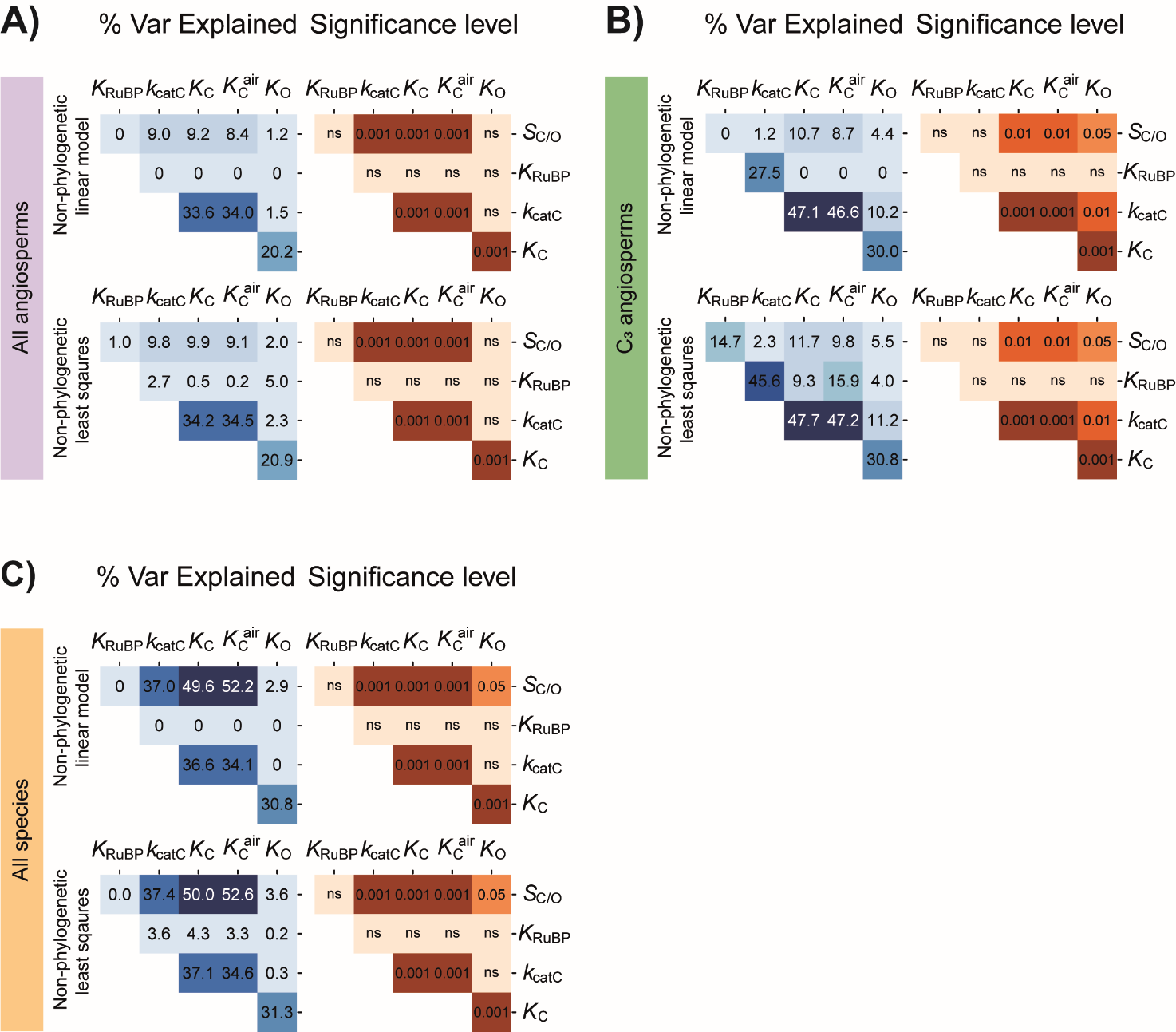

**Supplementary Figure S3.** **A)** Pairwise correlation coefficients (percent variance explained) and associated p-values between different rubisco kinetic traits assessed across the complete set of angiosperms using either non-phylogenetic linear models or least squares regression models. Significance values are represented as α levels, where; α = 0.001 if p < 0.001, α = 0.01 if 0.001 < p < 0.01, α = 0.05 if 0.01 < p < 0.05, and α = ns if p > 0.05. **B)** as in A but for C_3_ angiosperms only. **C)** as in A and B but for the complete set of photosynthetic organisms.

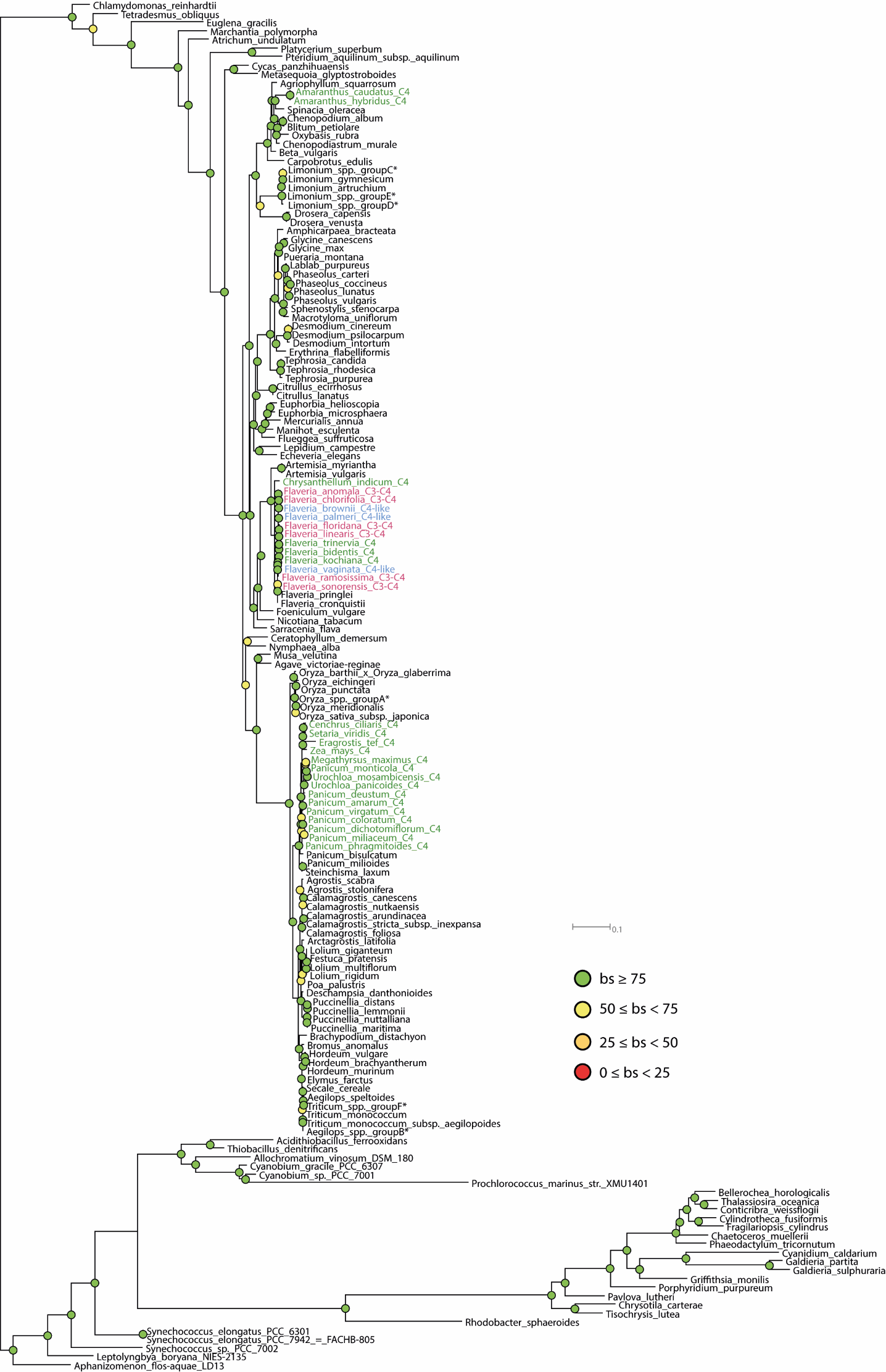

**Supplementary Figure S4.** Consensus phylogenetic tree of all photosynthetic organisms in this study inferred from the nucleotide sequence of the *rbcL* gene of the large subunit of the rubisco enzyme. The confidence of branch points on the tree are depicted as nodes which are colour coded by bootstrap support (bs) values. Where known, photosynthetic types other than the ancestral C_3_ pathway have been annotated on to species labels which are colour coded for visualisation, and include C_3_-C_4_ intermediates (red), C_4_-like (blue) and C_4_ plants (green). Species which shared identical sequences when considering the entire sequence length have been condensed into single nodes on the tree to avoid terminal zero-length branches and are signified by asterisks (see supplementary Figure S1 for further details). In addition to those species mentioned in supplementary Figure S1, an additional number of species exhibited terminal zero length branches relative to other nodes in this tree due to more stringent trimming of non-aligned positions at the terminal ends of the sequence alignment of all photosynthetic organisms. Of these additional species which exhibited terminal zero length branches, *Aegilops juvenalis* and *Aegilops vavilovii* were consensed into the node *Aegilops* spp. Group B, and *Triticum dicoccon, Secale cereale x Triticum aestivum*, *Triticum aestivum* and *Triticum timopheevii* were condensed into the node *Triticum* spp. Group F.

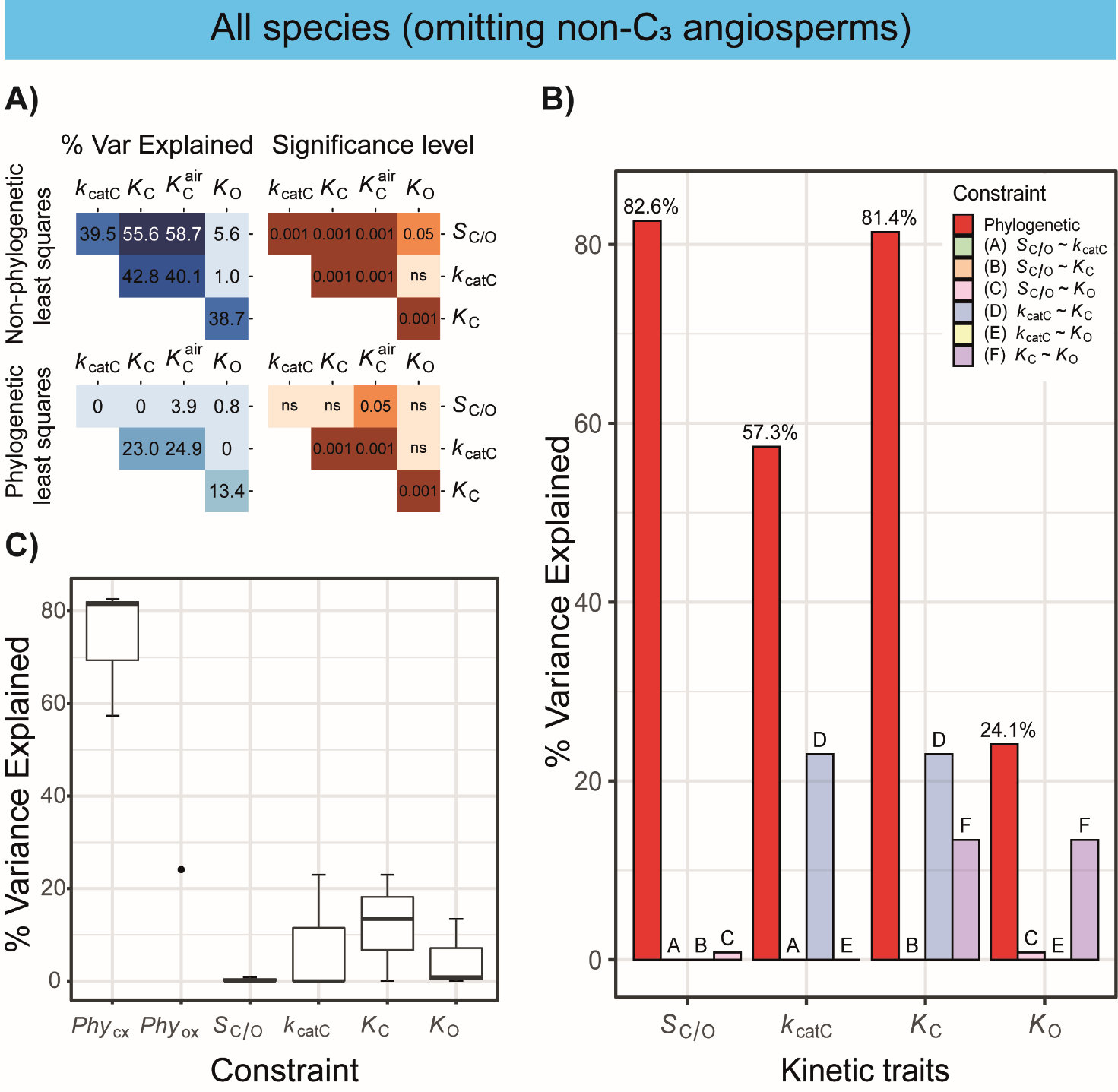

**Supplementary Figure S5.** Kinetic and phylogenetic constraints on rubisco adaptation across all studied photosynthetic organisms omitting known C_3_-C_4_, C_4_-like and C_4_ angiosperms **A)** Pairwise correlation coefficients (percent variance explained) and associated *p*-values between different rubisco kinetic traits assessed using non-phylogenetic least squares regression models or phylogenetic least squares regression models. Significance values are represented as α levels, where; α = 0.001 if *p* < 0.001, α = 0.01 if 0.001 < *p* < 0.01, α = 0.05 if 0.01 < *p* < 0.05, and α = ns if *p* > 0.05. **B)** The variation (%) in rubisco kinetic traits across photosynthetic organisms (omitting non-C_3_ angiosperms) that can be explained by phylogenetic constraint and each catalytic trade-off. **C)** Boxplot of all variation explained in each kinetic trait by kinetic trait correlations in comparison to variation explained by phylogeny in all photosynthetic organisms. The phylogenetic constraints on the carboxylase-related traits *Phy*_CX_ (includes *Phy_Sc/o_*, *Phy_Kcatc_*, and *Phy_Kc_*) and phylogenetic constraints on the oxygenase-related trait *Phy*_ox_ (includes *Phy_Ko_* only) are presented separately.

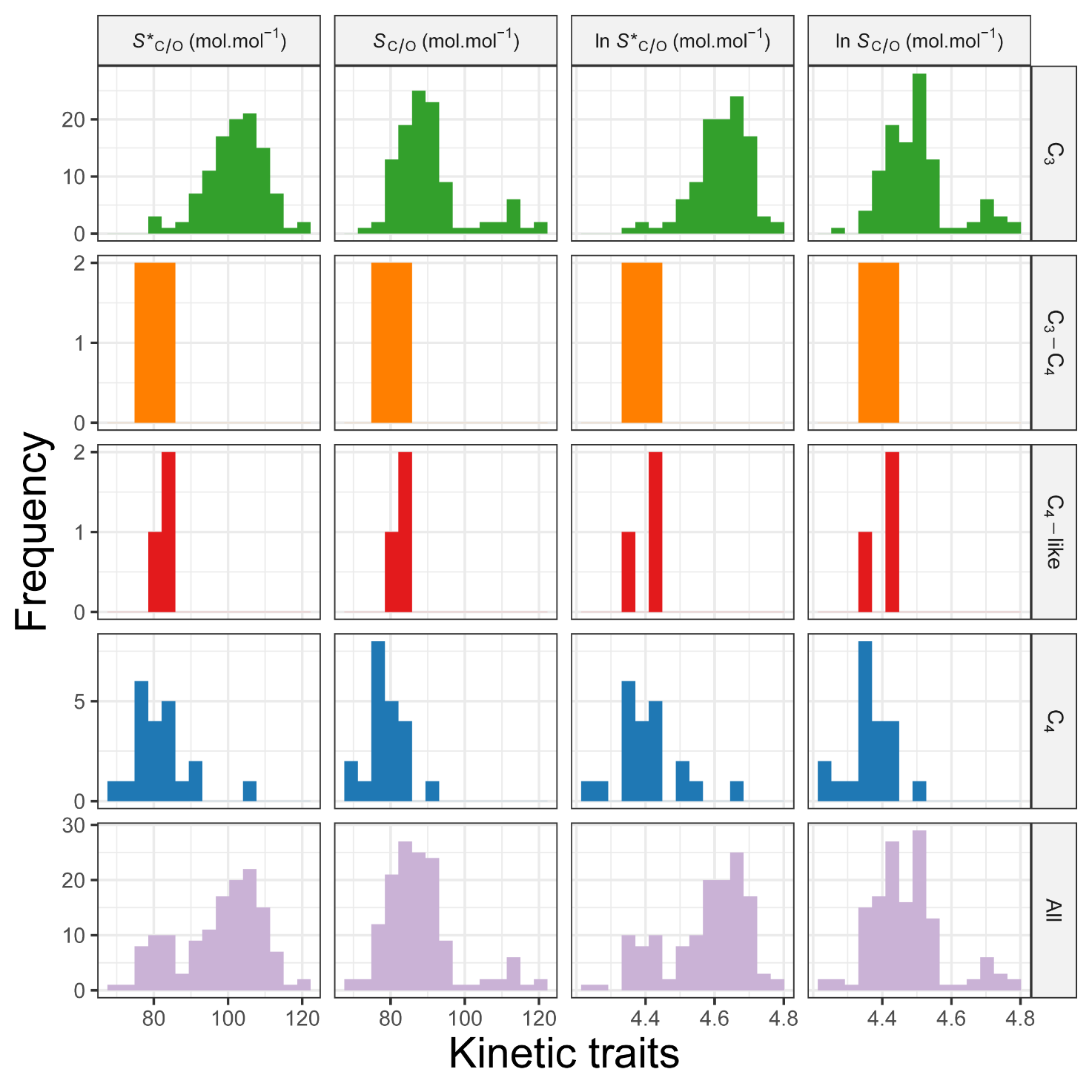

**Supplementary Figure S6.** The distributions of values for rubisco specificity in angiosperms before (*S**_C/O_) and after (*S*_C/O_) normalisation of measurements using the O_2_ electrode method (Orr *et al.*, 2016; Prins *et al.*, 2016) relative to those quantified using high precision gas-phase-controlled ^3^H-RuBP-fixation assays (Kane *et al.*, 1994). Species are grouped by their photosynthetic types (rows). Plants have been classified as those which perform C_3_ photosynthesis (C_3_; *n* = 107), C_4_ photosynthesis (C_4_; *n* = 6), C_3_-C_4_ intermediate (C_3_-C_4_; *n* = 3), C_4_-like (C_4_-like; *n* = 21). Both non-transformed and log-transformed non-normalised and normalised specificity values are shown, where respective units are shown in column labels.

**Supplementary Table S1.** The phylogenetic signal strength and associated significance level in rubisco kinetic traits across the phylogenetic tree in Supplementary Figure S2 in five signal detection methods. Statistics are rounded to 3 decimal places, and significance values are represented as α levels, where; α = 0.001 if *p* < 0.001, α = 0.01 if 0.001 < *p* < 0.01, α = 0.05 if 0.01 < *p* < 0.05, and α = ns if *p* > 0.05.

| **Kinetic Trait** | **C mean** | | **I** | | **K** | | **K *** | | **Lambda** | |
| --- | --- | --- | --- | --- | --- | --- | --- | --- | --- | --- |
|  | ***Stat*** | ***α*** | ***Stat*** | ***α*** | ***Stat*** | ***α*** | ***Stat*** | ***α*** | ***Stat*** | ***α*** |
| ***S*_C/O_** | 0.502 | 0.001 | 0.516 | 0.001 | 0.001 | 0.05 | 0.001 | 0.05 | 0.873 | 0.001 |
| ***k*_CatC_** | 0.343 | 0.001 | 0.303 | 0.001 | 0.002 | 0.001 | 0.003 | 0.001 | 0.940 | 0.05 |
| ***K*_C_** | 0.344 | 0.001 | 0.339 | 0.001 | 0.002 | 0.01 | 0.002 | 0.001 | 0.948 | 0.001 |
| ***K*_C_^air^** | 0.295 | 0.001 | 0.291 | 0.001 | 0.002 | 0.001 | 0.002 | 0.001 | 0.52 | 0.05 |
| ***K*_O_** | 0.125 | 0.05 | 0.142 | 0.05 | 0 | ns | 0 | ns | 0 | ns |

**Supplementary Table S2.** Significance level of assessed differences in rubisco kinetic traits between C_3_ and C_4_ angiosperms compared when measured in the absence of the phylogenetic tree, and when correctly accounting for the phylogenetic non-independence of species. Significance values are represented as α levels, where; α = 0.001 if *p* < 0.001, α = 0.01 if 0.001 < *p* < 0.01, α = 0.05 if 0.01 < *p* < 0.05, and α = ns if *p* > 0.05

| **Kinetic Trait** | ***α*** | |
| --- | --- | --- |
|  | – **Phylogenetic information** | **+ Phylogenetic information** |
| ***S*_C/O_** | 0.001 | 0.001 |
| ***k*_CatC_** | 0.001 | 0.001 |
| ***K*_C_** | 0.05 | 0.001 |
| ***K*_C_^air^** | 0.01 | 0.001 |
| ***K*_O_** | ns | ns |

**Supplementary Table S3.** The phylogenetic signal strength and associated significance level in rubisco kinetic traits across all studied photosynthetic organisms omitting known C_3_-C_4_, C_4_-like and C_4_ angiosperms using five different signal detection methods. Statistics are rounded to 3 decimal places and significance values are represented as α levels, where; α = 0.001 if *p* < 0.001, α = 0.01 if 0.001 < *p* < 0.01, α = 0.05 if 0.01 < *p* < 0.05, and α = ns if *p* > 0.05.

| **Kinetic Trait** | **C mean** | | **I** | | **K** | | **K *** | | **Lambda** | |
| --- | --- | --- | --- | --- | --- | --- | --- | --- | --- | --- |
|  | ***Stat*** | ***α*** | ***Stat*** | ***α*** | ***Stat*** | ***α*** | ***Stat*** | ***α*** | ***Stat*** | ***α*** |
| ***S*_C/O_** | 0.717 | 0.001 | 0.245 | 0.001 | 0.006 | 0.001 | 0.003 | 0.001 | 1.009 | 0.001 |
| ***k*_CatC_** | 0.53 | 0.001 | 0.313 | 0.001 | 0.002 | 0.001 | 0.001 | 0.01 | 0.955 | 0.001 |
| ***K*_C_** | 0.75 | 0.001 | 0.309 | 0.001 | 0.005 | 0.001 | 0.002 | 0.001 | 0.986 | 0.001 |
| ***K*_C_^air^** | 0.657 | 0.001 | 0.327 | 0.001 | 0.005 | 0.001 | 0.002 | 0.001 | 0.983 | 0.001 |
| ***K*_O_** | 0.332 | 0.001 | 0.151 | 0.01 | 0 | 0.05 | 0 | ns | 0.932 | 0.001 |

**Supplementary Table S4.** Key of abbreviated species labels as used in Figure 2 and Figure 3A.

| **Code** | **Species** |
| --- | --- |
| A.juv | *Aegilops juvenalis* |
| A.spe | *Aegilops speltoides* |
| A.vav | *Aegilops vavilovii* |
| A.squ | *Agriophyllum squarrosum* |
| A.sca | *Agrostis scabra* |
| A.sto | *Agrostis stolonifera* |
| A.cau | *Amaranthus caudatus* |
| A.hyb | *Amaranthus hybridus* |
| A.bra | *Amphicarpaea bracteata* |
| A.lat | *Arctagrostis latifolia* |
| A.myr | *Artemisia myriantha* |
| A.vul | *Artemisia vulgaris* |
| B.dis | *Brachypodium distachyon* |
| B.vul | *Beta vulgaris* |
| B.ano | *Bromus anomalus* |
| C.aru | *Calamagrostis arundinacea* |
| C.can | *Calamagrostis canescens* |
| C.fol | *Calamagrostis foliosa* |
| C.str | *Calamagrostis stricta subsp. inexpansa* |
| C.nut | *Calamagrostis nutkaensis* |
| C.cil | *Cenchrus ciliaris* |
| C.alb | *Chenopodium album* |
| C.mur | *Chenopodiastrum murale* |
| B.pet | *Blitum petiolare* |
| O.rub | *Oxybasis rubra* |
| C.ind | *Chrysanthellum indicum* |
| C.eci | *Citrullus ecirrhosus* |
| C.lan | *Citrullus lanatus* |
| D.dan | *Deschampsia danthonioides* |
| D.cin | *Desmodium cinereum* |
| D.int | *Desmodium intortum* |
| D.psi | *Desmodium psilocarpum* |
| E.far | *Elymus farctus* |
| E.tef | *Eragrostis tef* |
| E.fla | *Erythrina flabelliformis* |
| E.hel | *Euphorbia helioscopia* |
| E.mic | *Euphorbia microsphaera* |
| L.gig | *Lolium giganteum* |
| F.pra | *Festuca pratensis* |
| F.ano | *Flaveria anomala* |
| F.tri | *Flaveria trinervia* |
| F.bid | *Flaveria bidentis* |
| F.bro | *Flaveria brownii* |
| F.chl | *Flaveria chlorifolia* |
| F.cro | *Flaveria cronquistii* |
| F.flo | *Flaveria floridana* |
| F.koc | *Flaveria kochiana* |
| F.lin | *Flaveria linearis* |
| F.pal | *Flaveria palmeri* |
| F.pri | *Flaveria pringlei* |
| F.ram | *Flaveria ramosissima* |
| F.son | *Flaveria sonorensis* |
| F.vag | *Flaveria vaginata* |
| F.suf | *Flueggea suffruticosa* |
| F.vul | *Foeniculum vulgare* |
| G.can | *Glycine canescens* |
| G.max | *Glycine max* |
| H.vul | *Hordeum vulgare* |
| H.bra | *Hordeum brachyantherum* |
| H.mur | *Hordeum murinum* |
| L.pur | *Lablab purpureus* |
| L.cam | *Lepidium campestre* |
| L.artr | *Limonium artruchium* |
| L.gym | *Limonium gymnesicum* |
| L.mul | *Lolium multiflorum* |
| L.rig | *Lolium rigidum* |
| M.uni | *Macrotyloma uniflorum* |
| M.esc | *Manihot esculenta* |
| M.max | *Megathyrsus maximus* |
| M.ann | *Mercurialis annua* |
| M.vel | *Musa velutina* |
| N.tab | *Nicotiana tabacum* |
| xO.hyb | *Oryza barthii x Oryza glaberrima* |
| O.eic | *Oryza eichingeri* |
| O.mer | *Oryza meridionalis* |
| O.pun | *Oryza punctata* |
| O.sat | *Oryza sativa* subsp. *japonica* |
| P.ama | *Panicum amarum* |
| P.bis | *Panicum bisulcatum* |
| P.col | *Panicum coloratum* |
| P.deu | *Panicum deustum* |
| P.dic | *Panicum dichotomiflorum* |
| P.mid | *Panicum milioides* |
| P.mic | *panicum miliaceum* |
| P.mol | *Panicum monticola* |
| P.phr | *Panicum phragmitoides* |
| P.vir | *Panicum virgatum* |
| P.car | *Phaseolus carteri* |
| P.coc | *Phaseolus coccineus* |
| P.lun | *Phaseolus lunatus* |
| P.vul | *Phaseolus vulgaris* |
| P.pal | *Poa palustris* |
| P.dis | *Puccinellia distans* |
| P.lem | *Puccinellia lemmonii* |
| P.mar | *Puccinellia maritima* |
| P.nut | *Puccinellia nuttalliana* |
| P.mon | *Pueraria montana* |
| S.cer | *Secale cereale* |
| S.vir | *Setaria viridis* |
| S.ste | *Sphenostylis stenocarpa* |
| S.ole | *Spinacia oleracea* |
| S.lax | *Steinchisma laxum* |
| T.dic | *Triticum dicoccon* |
| T.mon | *Triticum monococcum* |
| T.tim | *Triticum timopheevii* |
| T.can | *Tephrosia candida* |
| T.pur | *Tephrosia purpurea* |
| T.rho | *Tephrosia rhodesica* |
| xTrit | *Secale cereale x Triticum aestivum* |
| T.aes | *Triticum aestivum* |
| T.aeg | *Triticum monococcum subsp. aegilopoides* |
| U.mos | *Urochloa mosambicensis* |
| U.pan | *Urochloa panicoides* |
| Z.may | *Zea mays* |
| O.sppA* | *Oryza* spp. Average Group A |
| A.sppB* | *Aegilops* Spp. Average Group B |
| L.sppC* | *Limonium* Spp. Average Group C |
| L.sppD* | *Limonium* Spp. Average Group D |
| L.sppE* | *Limonium* Spp. Average Group E |

* Means have been taken between species which share identical *rbcL* sequences. *Oryza* spp. Average Group A include *O. longistaminata*, *O. glaberrima*, *O. sativa* f. *spontanea*, *O. sativa* subpsp. *indica* and *O. glumipatula*; *Aegilops* spp. Average Group B include *A. triuncialis*, *A. uniaristata*, *A. tauschii*, *A. comosa*, *A.* *biuncialis*, *A. cylindrica*; *Limonium* spp. Average Group C include *L. antonii-llorensii*, *L. gibertii*, *L. biflorum*; *Limonium* spp. Average Group D include *L.* *echioides*, *L. barceloi*, *L.* *balearicum*, *L. companyonis*; and *Limonium* spp. Average Group E include *L. ejulabilis*, *L. retusum*, *L. leonardi-llorensii*, *L. magallufianum*, *L. grosii*.
